# Goldilocks conundrum explains cryoinjury in slow-cooled amphibian embryonic cells

**DOI:** 10.1101/2025.11.24.690295

**Authors:** Roshan Patel, Rose Upton, Simon Clulow, Brett Nixon, Michael Mahony, John Clulow

**Author notes:** **Correspondence should be addressed to Roshan Patel:**.

## Abstract

Cryopreservation of intact fish and amphibian eggs and embryos is challenging due to the sheer size and yolk content, which prevents proper dehydration and causes lethal intracellular ice formation during cooling. Alternatively, cryopreservation of dissociated embryonic cells allows to biobank diploid genomes. However, amphibian and fish embryos have cells of varying sizes throughout the developing embryo and cell size distribution remains understudied in cryopreservation. This study examined cell size effects during cryopreservation of dissociated embryonic cells from two amphibian species. Most work used *Limnodynatses peronii* blastula, gastrula, and neurula cells cryopreserved with 10% dimethyl sulfoxide (DMSO) and sucrose at 0%, 1%, or 10%. Increasing sucrose concentration improved post-thaw recovery of membrane intact cells and cell concentrations, with gastrula and neurula cells having better recovery than blastula cells. An interaction amongst cryoprotectant concentration, cell size, and embryonic stage was observed. Higher sucrose improved recovery of larger cells, but reduced recovery of smaller cells. These results supported a “Goldilocks” model of cryoinjury, in which larger cells require more time to dehydrate adequately, while smaller cells are damaged by excessive dehydration and solute effects, implying that optimal cryoprotectant conditions differ by cell size. Based on post-thaw recovery, 10% DMSO + 10% sucrose was optimal and successfully applied to the cryopreservation of neurula cells from the threatened *Rawlinsonia littlejohni*, marking the first report of embryonic cell cryopreservation in a threatened amphibian. Our results demonstrated the importance of cell size effects on cryoinjury and their consideration in the application of cryopreservation for amphibian conservation.

In brief: Since intact amphibian eggs and embryos cannot be cryopreserved, dissociated embryonic cell cryopreservation provides an alternative avenue for biobanking embryonic diploid genomes; however, cell size effects on cryoinjury is poorly understood and the developing amphibian embryo consists of different sized cells. We demonstrated that cell size influences cryopreservation outcomes by cryopreserving embryonic cells from three developmental stages under varying cryoprotectant concentrations.

## Introduction

Cryopreservation of eggs and embryos of fish and amphibians (anamniotes) has proven a great challenge to cryobiology (Clulow et al., 2022; Strussmann et al., 1999). This is unfortunate because many species in both groups are endangered (IUCN, 2023), and the cryopreservation of eggs and embryos would benefit assisted reproductive technologies (ART) and biobanking for their conservation (Della Togna et al., 2020; Kusuda et al., 2004; Strussmann et al., 1999). However, technological limitations for egg and embryo cryopreservation have restricted the application of ARTs and biobanking for threatened anamniotes (Clulow et al., 2022; Strussmann et al., 1999). Anamniote eggs and embryos are large and yolk-rich, which limits cryoprotectant permeability, restricts dehydration, and favours lethal intracellular ice formation during cryopreservation (Clulow et al., 2022; Hagedorn et al., 1997, 1998, 2004). The size of anamniote eggs and embryos is a consequence of the evolution of lecithotrophic nutrition and reproductive mode (Blackburn, 2000, 2015). By comparison, mammalian oocytes and embryos, which are smaller (80-150 µm in diameter) have been successfully cryopreserved (Clulow et al., 2019).

Despite recent success of vitrified embryos in the zebrafish, which are small in diameter (Khosla et al., 2020), most reports of diploid genome preservation of lecithal embryos have utilised slow cooling of embryonic cells. This approach has generated viable chimeras in some fish species (Calvi & Maisse, 1998; Kawakami et al., 2010; Kusuda et al., 2004; Yoshizaki & Lee, 2018). In amphibians, only one study reported recovery of embryonic cells by slow cooling penetrating cryoprotectants (Lawson et al., 2013). Cryopreservation of embryonic cells by slow cooling is likely to continue to be the main form of diploid embryo genome cryopreservation for most aquatic species. However, few studies of the cryopreservation of embryonic cells from lecithotrophic species have investigated the mechanisms of cryoinjury essential for optimizing such protocols (Strussmann et al., 1999).

Cell size is known factor affecting cell response and recovery during cryopreservation (Fadda et al., 2010; Fadda et al., 2011), but poorly studied. This is probably due to the little variability in cell size distribution in many cryopreservation systems (Fadda et al., 2010). Aquatic lecithotrophic embryos show a vast variation in cell size within and between developmental stages—from early cleavage to neurulation—with cell volume decreasing by up to 10 -fold during cleavage (species in this study).

In this study, we demonstrated the importance of cell size distribution as a predictor of cryoinjury and contributor to post-thaw variation in viable embryonic cell recovery in two amphibian species (*Limnodynastes peronii* and *Rawlinsonia littlejohni, formerly Litoria littlejohni*) (Donnellan et al., 2025)), the latter being highly threatened. The main experiment involved cryopreserving *L. peronii* embryonic cells from three developmental stages using three levels of sucrose followed by applying the optimal cryoprotectant combination to cryopreserve embryonic cells from *R. littlejohni*. From these experiments, we developed an integrated conceptual model describing the mechanism of cryoinjury in early embryonic cells.

## Materials and Methods

### Animal Husbandry

#### Limnodynastes peronii

Male and female *L. peronii* were collected from Kooragang Island and the Watagan Mountains,NSW, Australia, and housed at the University of Newcastle. Individuals were housed in plastic terraria (30 × 18 × 20 cm) containing 75% terrestrial (autoclaved gravel, leaf litter, and artificial plants) and 25% aquatic (aged tap water) environments, with three frogs per terrarium (one male and two females). Frogs were fed live crickets dusted with calcium and vitamin powder (Multical Dust; Vetafarm, Wagga Wagga, NSW, Australia) and water was changed twice a week. All individuals collected from the wild underwent a heat treatment protocol to eliminate potential chytridiomycosis, using UV- and temperature-controlled cabinets (TRISL–1175, Thermoline Scientific Equipment, Wetherill Park, NSW) on a 12:12 UV/daylight cycle starting at 24°C. Temperature was increased by 2°C per day to 36 °C, then raised to 37°C for six hours, a temperature known to kill *Batrachochytrium dendrobatidis* within four hours (Johnson et al., 2003). The temperature was then reduced by 2°C per day back to 24°C, after which terraria were maintained at room temperature on a 12:12 light/dark cycle.

#### Rawlinsonia littlejohni

Adult male and female *R. littlejohni* were collected in the Watagans Mountains, NSW and housed in glass terraria (60 x 50 x 30 cm) with autoclaved sand, leaflitter, and artificial plants, with a water dish containing 10% Simplified Amphibian Ringer (SAR; 113 mM NaCl, 1 mM CaCl_2_, 2 mM KCl, 3.6 mM NaHCO_3_; ∼220 mOsm/kg; (Browne et al., 1998). *R. littlejohni*, a winter-breeding species, were maintained at 15-19 °C. Individuals were fed dusted live crickets and water changed weekly. All individuals received voriconazole prophylactically to eliminate chytrid infection. All experiments were carried out in accordance the University of Newcastle’s animal ethics approval, A-2022-207. Animals were collected and held under NSW Scientific Licence SL101269.

#### Embryo collection

Three L. peronii spawns were collected following natural spawning events in terraria. Spawns were transferred to 150 × 15 mm Petri dishes containing 10% Niu-Twitty solution (McKinnell, 1978) (5.8 mM NaCl, 0.07 mM KCl, 0.04 mM Ca(NO₃)₂, 0.04 mM MgSO₄, 0.03 mM Na₂HPO₄, 0.015 mM KH₂PO₄, 0.24 mM NaHCO₃) prepared in reverse osmosis water and held at 13°C in a temperature- and humidity-controlled cabinet (Thermoline Scientific, Australia; TRH-300-GD) to slow development. Embryos were sampled at Gosner stages 9 (blastula), 11 (gastrula), and 13 (neurula) (Gosner, 1960).

#### Preparation of isolated embryonic cells

Thirty *L. peronii* embryos were removed from each spawn at blastula, gastrula, and neurula (Gosner, 1960). Foam and egg jelly were removed from embryos by rolling them across flattened paper towel. Embryos were then dissociated in 2 mL of 2.5% Bovine Serum Albumin (BSA) (Sigma-Aldrich A7906-50G) prepared in modified Niu-Twitty (50.36 mM NaCl, 0.67 mM KCl, 3.65 mM Na_2_HPO_4_, 0.85 mM KH_2_PO_4_, 2.38 mM NaHCO_3_) (McKinnell, 1978) via dissection with forceps. The dissected embryonic cells were pipetted into 2mL Eppendorf tubes and placed on ice.

#### Cell counts, cell viability, and photo acquisition

Embryonic cells were assessed in a Neubauer hemacytometer under a Nikon Eclipse e200 compound microscope at 400x magnification for cell concentration and membrane integrity. Cells (10 µL per chamber) were counted in all four primary squares. Membrane integrity was determined using a 0.5% (w/v) trypan blue exclusion assay (1:1 cell-to-dye ratio), with unstained cells scored as viable and stained cells as non-viable (Figure 1). At least 100 cells were counted per sample, with two replicate counts per measurement. Images (10 per replicate per developmental stage pre-freeze) were captured using a Tucsen ISH500 camera on an Olympus CX41 microscope at 200× magnification, and cell diameters were measured in TCapture software (v4.3.0.605). A total of 90 images were obtained pre-freeze across all replicates and stages.

**Figure 1:**
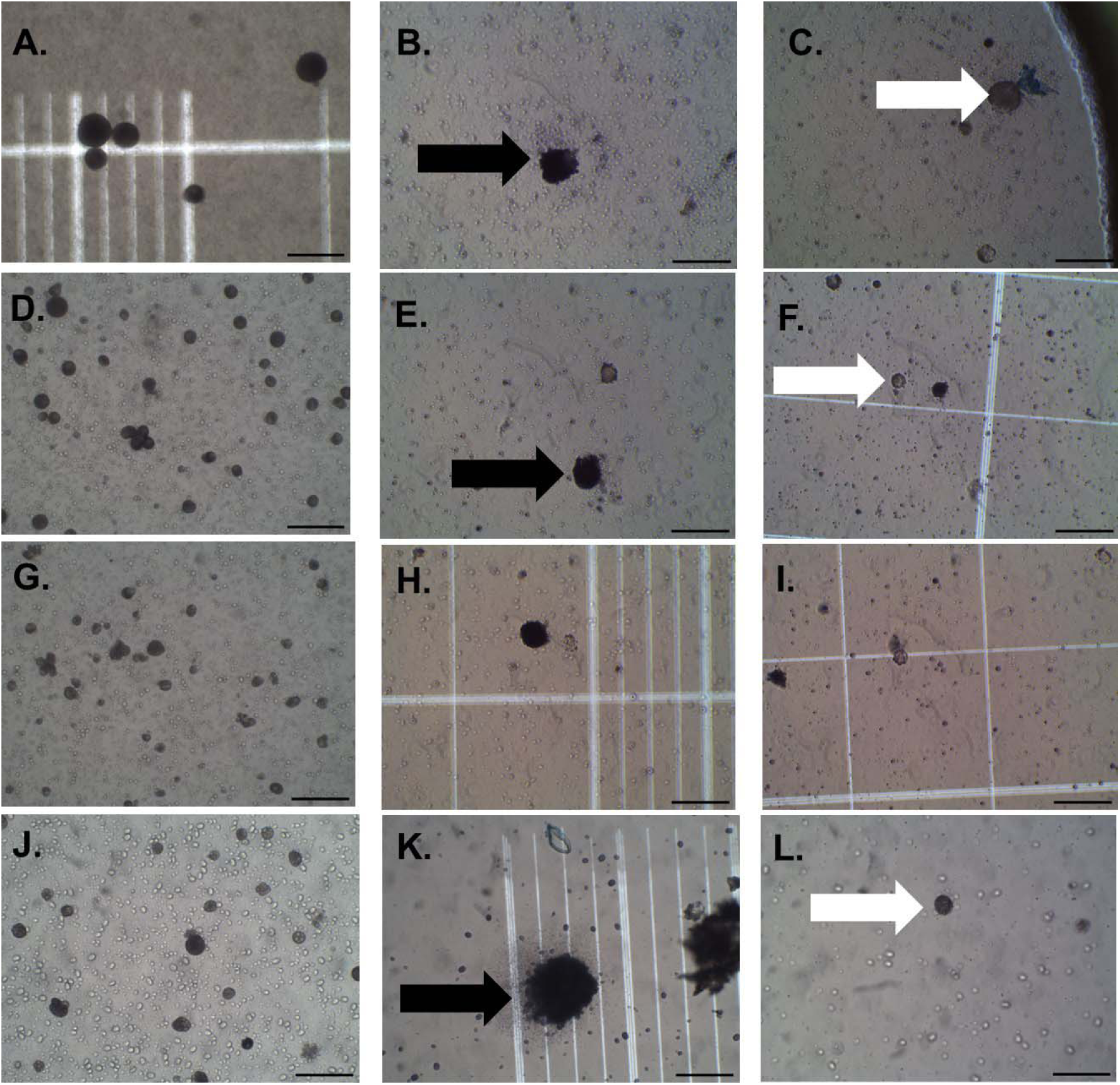
Pre-freeze (1A) and post-thaw blastula (1B, 1C), pre-freeze (1D) and post-thaw gastrula (1E, 1F), and pre-freeze (1G), post-thaw neurula (1H, 1I) images from *L. peronii*, and pre-freeze (1J) and post-thaw (1K and 1L) neurula from *R. littlejohni*. Post-thaw *L. peronii* blastula (1B, 1C), gastrula (1E, 1F), neurula (1H, 1I), and *R. littlejohni* neurula (1K and 1L) have been treated with trypan blue. Black arrows point to stained cells, and white arrows point to unstained cells. Scale bar = 100 µm (added with ImageJ).

#### Cryopreservation

Three cryoprotectant solutions (prepared in modified Niu-Twitty solution) were used: 10% v/v DMSO (Ajax Finechem, AJA2225; 10D0S), 10% v/v DMSO + 1% w/v sucrose (AJA530; 10D1S), and 10% v/v DMSO + 10% w/v sucrose (10D10S). Prior to cryopreservation, 250 µL of cryoprotectant was added to 50 µL of embryonic cell suspension in Eppendorf tubes on ice. A total of 300 µL of each mixture was loaded into 500 µL straws (Minitube Eco Straw, Ref. 13408/3010) and sealed with polyvinyl alcohol powder. Five straws were prepared for each combination of spawn, developmental stage, and cryoprotectant. Straws were frozen in a programmable freezer (Cryologic CL-3300, Australia) following protocol: held at 10°C for ten min, cooled to −7°C at −1 °C/min, held for ten min, cooled to −30°C at −1°C/min, held for ten min, then cooled by free fall to −80°C (Lawson et al. 2013). Straws were then plunged into liquid nitrogen and stored in a dewar for two weeks before post-thaw assessment.

#### Post-thaw analysis

Straws were thawed at room temperature (∼21°C) on the laboratory bench and the contents emptied into a 1.5 mL Eppendorf tube on ice. Post-thaw cell concentration, membrane integrity, and diameter were assessed as described above. Due to low recovery in some treatments, counts of 100 cells per sample were not always possible. At least 10 images per straw were acquired, totaling 1,604 images across all replicates.

### Application to R. littlejohni

Neurula cells of *R. littlejohni* were cryopreserved using 10D10S following the same protocol as for *L. peronii*. Cells were obtained from three IVF combinations (sperm from three males, eggs from one female), using abnormally developing embryos while normal embryos were left to develop. Eggs were hormonally induced via pituitary extracts (one male *Limnodynastes dumerilii*, one female *Litoria aurea*, two male *L. peronii*, and two male *Litoria caerulea* pituitaries in SAR) injected into the dorsal lymph sac, with oviposition 24 h later. Sperm for IVF was collected after induction with human chorionic gonadotrophin (5 IU/g body weight in SAR).

Embryos were dejellied in 1% w/v cysteine (Sigma-Aldrich c1276) (prepared in modified Niu- Twitty and pH=8) for 17 min. Five embryos from the three IVF combinations were dissociated (one in 500 µL 2.5% BSA) and kept on ice. Pre-freeze assessment, cryopreservation (three straws per embryo) and post-thaw analysis followed the *L. peronii* protocol. Pre- and post-thaw imaging yielded 54 and 181 photos, respectively.

### Data analysis

#### L. peronii

Generalised Linear Mixed Models (GLMMs) were used to analyse the effects of embryonic stage and cryoprotectant on cell membrane integrity (binary logistic regression; weights = total cell counts), with overdispersion addressed via an observation-level random effect (OLRE). Cell concentration (cells/mL) pre- and post-thaw was analysed using GLMMs with a negative binomial distribution; raw counts were adjusted to cells/mL using an offset based on dilution, volume, and quadrats counted. Embryonic stage and cryoprotectant were fixed effects, with spawn ID as a random effect.

Pre- and post-thaw cell size distributions across stages and cryoprotectants were compared using Kolmogorov-Smirnov tests and visualised with histograms showing means and medians. To assess effects on size-class distributions, GLMMs with binary logistic regression (interpreted as the proportion counts per size class) were used (weights = total cell counts per straw), with interaction of cryoprotectant, embryonic stage, and size class as the fixed effect and OLRE for overdispersion. Cells were grouped into 10 µm size classes (0–10 µm to 111–120 µm).

#### R. littlejohni

Analyses followed the same approach as for *L. peronii*. GLMMs were used for membrane integrity, cell concentration, and size-class recovery. Cell size distributions were compared using Kolmogorov-Smirnov tests and GLMMs with interaction effect of cryoprotectant and size class. Cells were grouped into 10 µm size classes from 0–10 µm to 141–150 µm.

Data analysis was performed using the R programming language (Version 4.1.2) in RStudio with the following packages: glmmTMB for negative binomials for cell concentrations (cell/mL) and lme4 other general linear mixed modelling (GLMM) for membrane integrity and cell size count distributions (Bates, 2014; Brooks et al., 2017); emmeans was used to model Estimated Marginal Means (EMM), calculate 95% confidence intervals (CI), and odds ratios (OR) for membrane integrity, cell concentration, and cell size count distributions (Lenth, 2022); DHARMa was used to assess QQplot residuals, distribution, and dispersion (Hartig, 2018); gridExtra, ggplot2 and ggeffects were used for plotting (Auguie & Antonov, 2017; Wickham et al., 2019, 2024).

## Results

### Cryopreservation associated with loss of cells and reduced viability in recovered cells for L. peronii

There was a significant effect of sucrose (likelihood ratio test (LRT) χ^2^(3) = 98.51, P-value: 3.24X10^-21^) and embryonic stage (LRT χ^2^(2) = 372.56, P: 1.26X10^-81^) on cell concentration post-cryopreservation with more cells being recovered as the sucrose concentration increased (Fig. 2A). Odds ratios (OR) and 95% confidence intervals (CI) were calculated to compare treatments. Post-thaw cell concentrations for blastula, gastrula, and neurula (72,717, 373,980, and 639,316 cells/mL, respectively) cells cryopreserved with 10D10S were significantly higher than 10D0S (OR: 1.88, 95% CI: 1.623-2.178) and 10D1S (OR: 1.509-fold,95% CI 1.308-1.741). Post-thaw cell concentrations (cells/mL) for blastula, gastrula, and neurula cells were 38,683, 198,944, and 340,093 for 10D0S, and 48,187, 247,823, and 423,651 for 10D1S, respectively.

**Figure 2.**
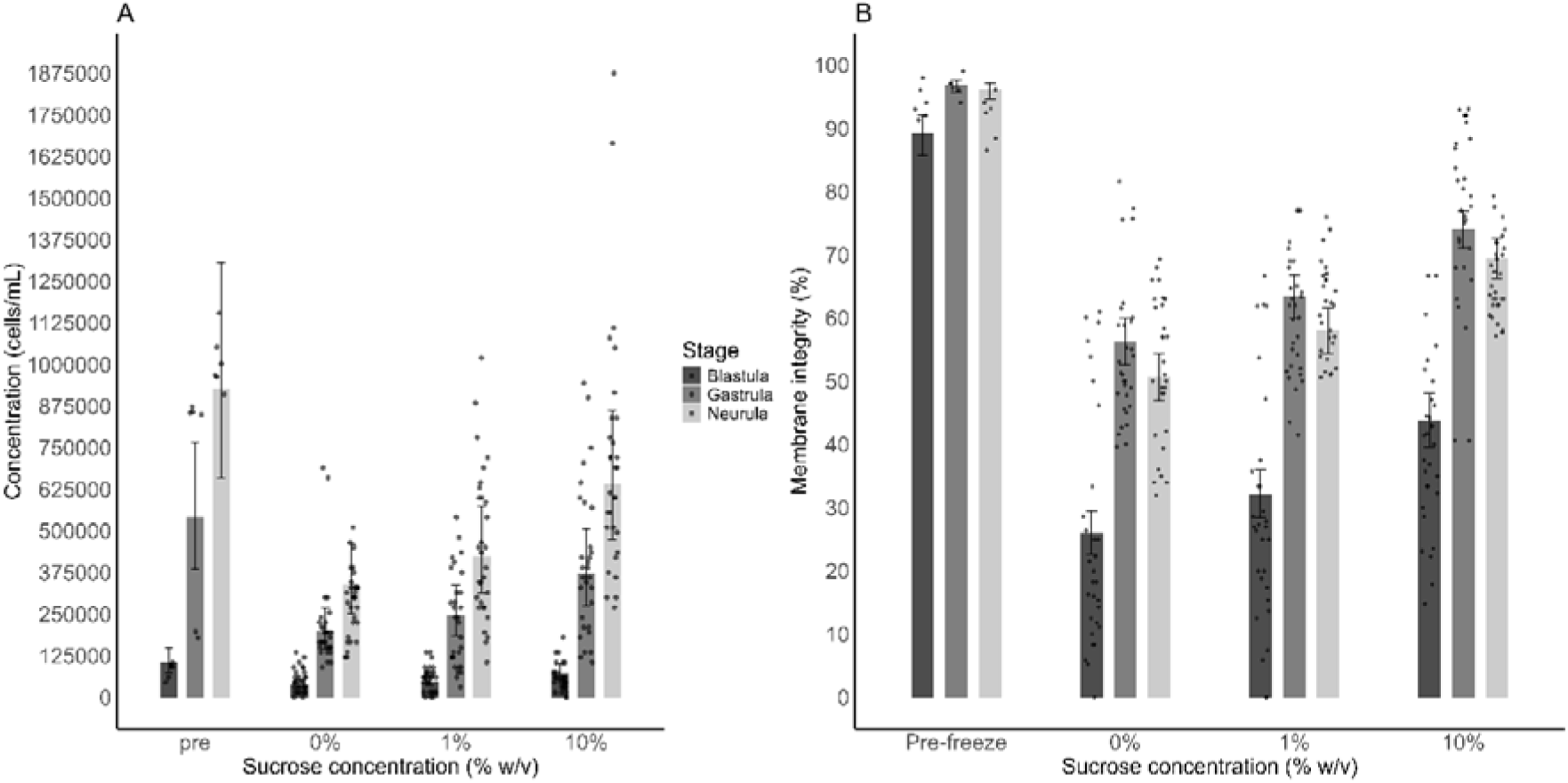
Cell concentration and cell membrane integrity after of *L. peronii* embryonic cells cryopreserved with different sucrose concentrations (10% v/v DMSO with either 0, 1, or 10% w/v sucrose). 2A. Pre-freeze and post-thaw cell concentrations (cells/mL) of dissociated embryos from blastula, gastrula, and neurula stages. GLMM negative binomial. 2B. Percentage of membrane intact cells post-thaw in blastula, gastrula, and neurula cells. GLMM binomial. EMMs were plotted along with 95% CI and raw data points (black dots) for 2A and 2B.

There was a significant effect of sucrose concentration (LRT χ^2^(3) = 263.72, P: 7.06X10^-57^) and embryonic cell stage and LRT χ^2^(2) = 180.73, P-value: 5.69X10^-40^) on cell membrane integrity post-thaw (Fig. 2B). Odds ratios (OR) and 95% confidence intervals (CI) were calculated to compare treatments. As sucrose concentration increased, so did the percentage of membrane intact cells. Post-thaw membrane intact cell percentages for blastula, gastrula, and neurula (43.7%, 74.1%, and 69.5%) cells cryopreserved with 10D10S were significantly higher than with 10D0S (OR: 2.223, 95% CI: 1.871-2.641) and 10D1S sucrose (OR: 1.647, 95% CI: 1.387-1.957). Post-thaw membrane intact cell percentages for blastula, gastrula, and neurula were 25.9%, 56.2%, and 50.6% with 10D0S, and 32.1%, 63.4%, and 58% with 10D1S, respectively. For 10D10S, gastrula and neurula cells had a significantly higher percentage of membrane intact cells recovered post thaw than blastula cells (OR: 3.675; 2.93, 95% CI: 3.062-4.411; 2.448-3.506, respectively).

Cryopreservation had a significant effect on the cell size distribution of cells post-thaw compared to pre-freeze in all embryo stages and all cryoprotectant combinations (Figs 3-6). There was a significant interaction amongst cryoprotectant concentration, size class, and embryonic stage (LRT χ2(30) = 136.21, P: 1.76X10^-15^) on cell size class distributions pre- freeze and post-thaw.

**Figure 3:**
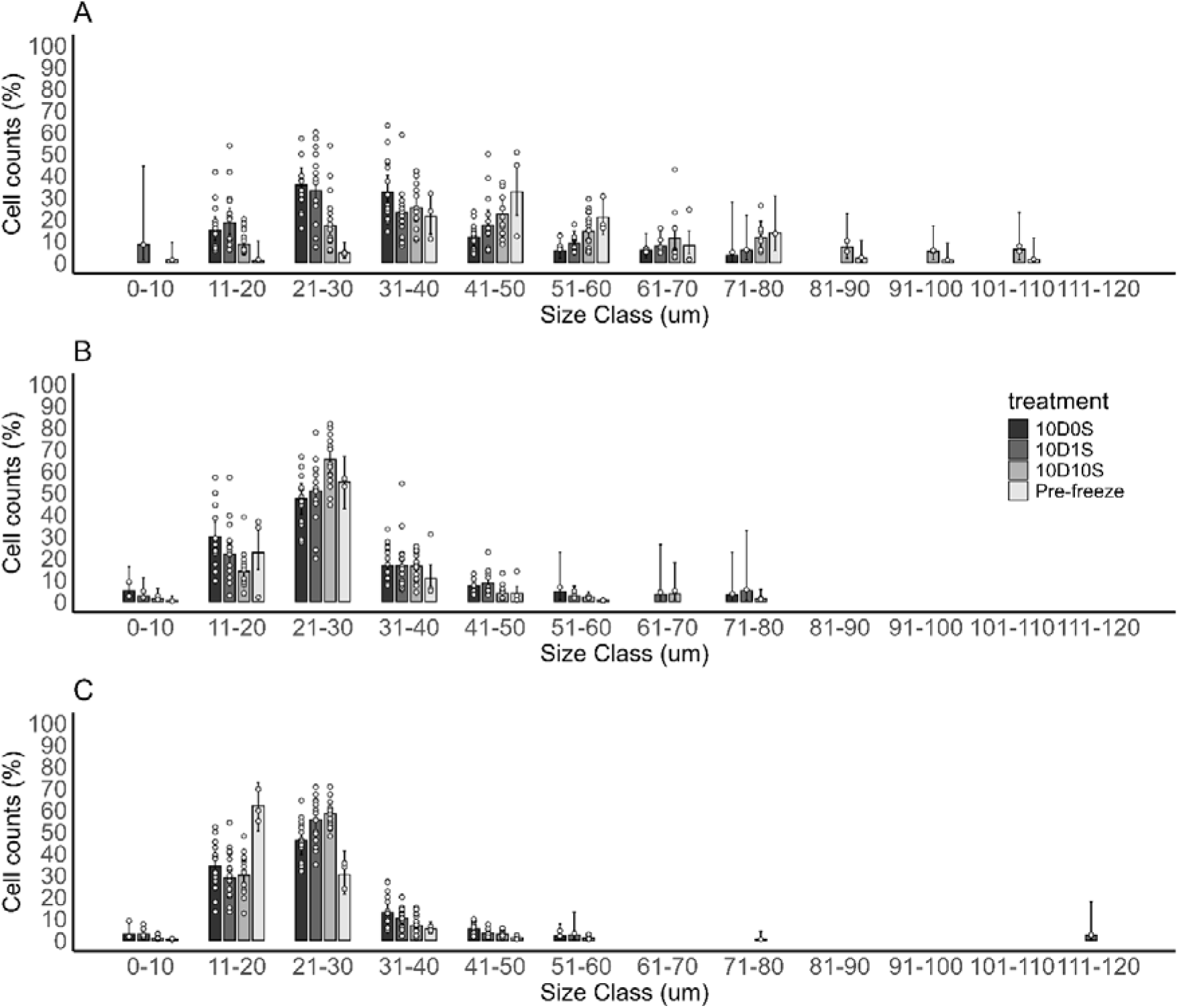
*L. peronii* cell diameter class distributions pre-freeze and post-thaw (all cryoprotective additives) for blastula (A), gastrula (B), and neurula (C). GLMM binomials. EMMs for the percentage of cells in each cell size class along with 95% CI and raw data (white dots).

### Blastula

Cell size distributions for *L. peronii* blastulas post-thaw differed significantly from pre-freeze for all cryoprotectants: 10D0S (D = 0.62, P = 2.51×10⁻³□), 10D1S (D = 0.54, P = 2.44×10⁻³□), and 10D10S (D = 0.23, P = 1.35×10⁻□). Fewer large cells were present post-thaw, indicating greater loss of larger blastula cells (Fig. 4A–C). The loss of large cells in lower CPA treatments (10D0S and 10D1S) corresponded with a significantly increased proportion of smaller (11–20 µm) cells post-thaw—10D0S (EMM: 14.9%, OR: 0.05, 95% CI: 0.004–0.65) and 10D1S (EMM: 18.4%, OR: 0.04, 95% CI: 0.003–0.50)—relative to pre-freeze (0.86%) (Figure 5A). Conversely, 10D10S showed significantly fewer small cells (11– 20 µm; EMM: 8.5%) than 10D0S (OR: 0.53, 95% CI: 0.29–0.99) and 10D1S (OR: 0.41, 95% CI: 0.23–0.75). Its protective effect was evident in the recovery of larger cells (81–90 µm; 7.3% post-thaw vs. 2.1% pre-freeze).

**Figure 4:**
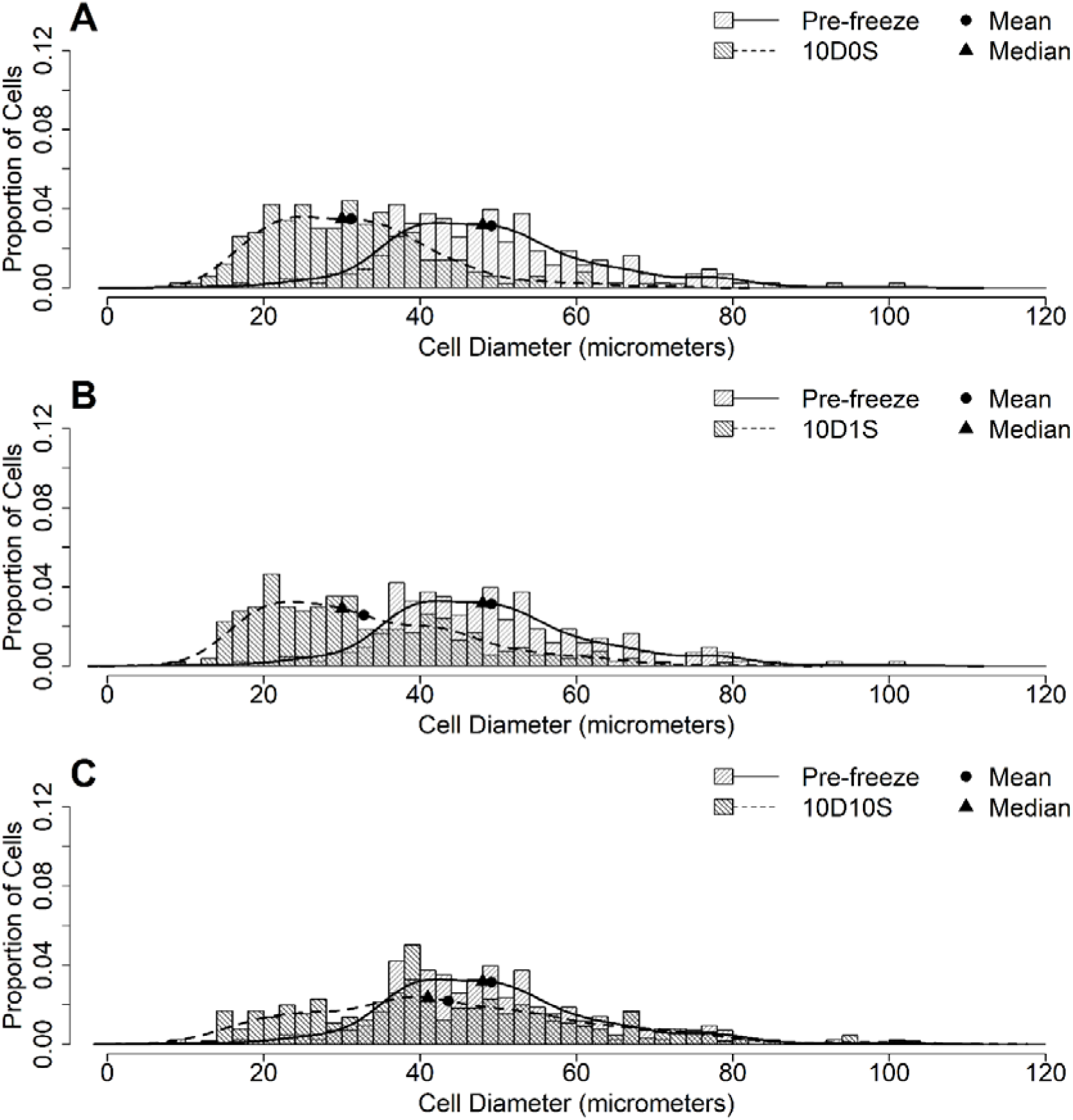
Cell size distributions of *L. peronii* blastulas pre-freeze (mean 49.13 µm, median 48 µm) and post-thaw in (A) 10D0S (mean 31.18 µm, median 30 µm), (B) 10D1S (mean 32.8 µm, median 30 µm), and (C) 10D10S (mean 43.59 µm, median 41 µm).

**Figure 5:**
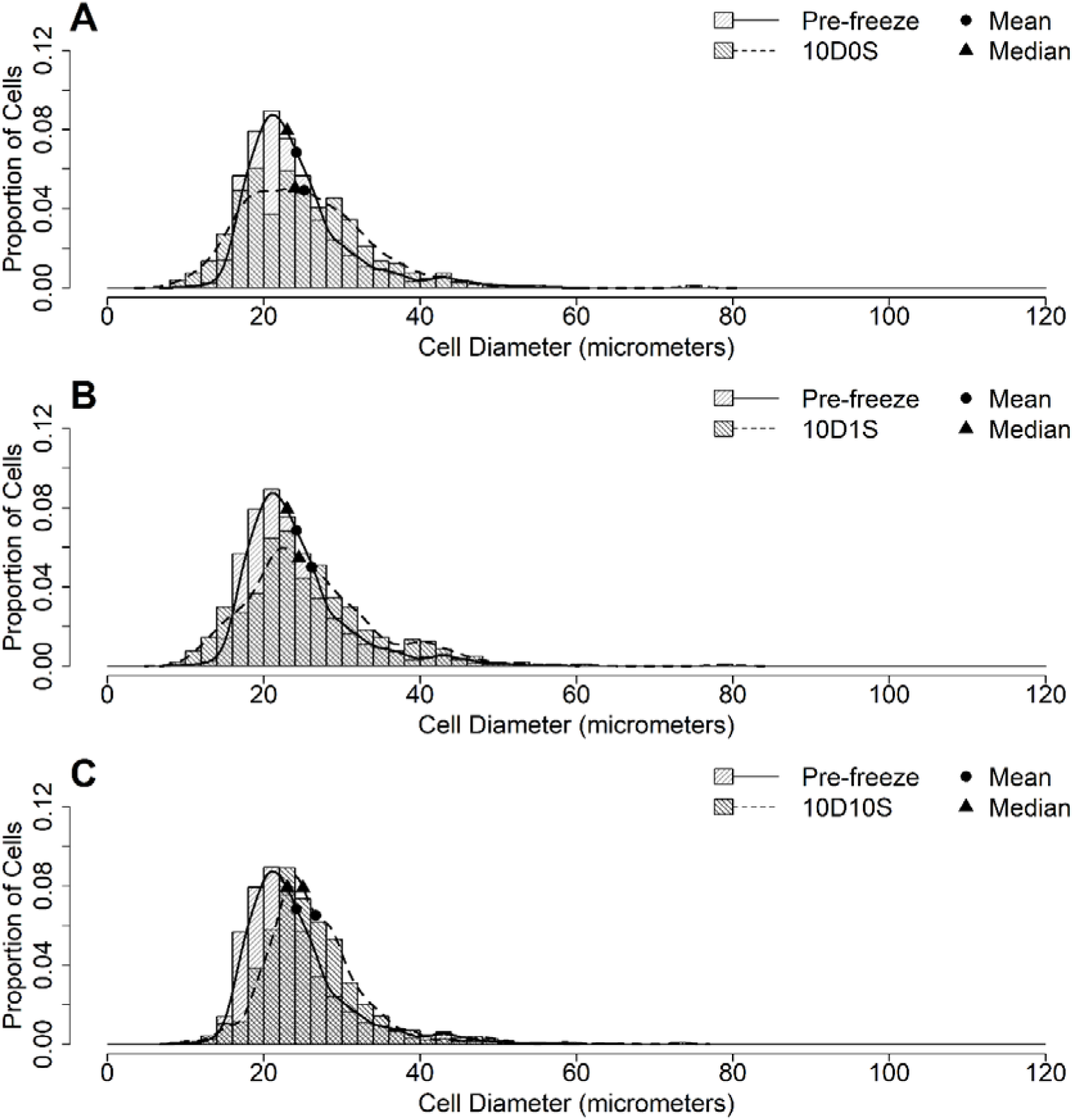
Cell size distributions of *L. peronii* gastrulas pre-freeze (mean 24.17 µm, median 23 µm) and post-thaw in (A) 10D0S (mean 25.18 µm, median 24 µm), (B) 10D1S (mean 26.12 µm, median 24.5 µm), and (C) 10D10S (mean 26.65 µm, median 25 µm).

### Gastrula and Neurula

Post-thaw cell size distributions showed a higher frequency of larger cells across all cryoprotectants for both gastrula and neurula stages (Figs.5-6): 10D0S (gastrula D = 0.14, P = 1.1×10⁻□; neurula D = 0.27, P = 6.49×10⁻²□), 10D1S (gastrula D = 0.16, P = 4.9×10⁻□; neurula D = 0.32, P = 2.15×10⁻□¹), and 10D10S (gastrula D = 0.24, P = 4.66×10⁻¹□; neurula D = 0.32, P = 2.1×10⁻□).

### Gastrula

For gastrulas (Fig. 3B), cell counts in the 11–20 µm class significantly declined with increasing sucrose concentration—from 29.99% (10D0S) to 21.95% (10D1S) and 14.01% (10D10S)—despite no overall difference from pre-freeze (22.62%) (10D10S vs. 10D0S: OR = 0.38, 95% CI = 0.25–0.58; 10D10S vs. 10D1S: OR = 0.58, 95% CI = 0.38–0.88). In contrast, cell counts in the 21–30 µm class significantly increased with sucrose—from 47.22% (10D0S) to 51.02% (10D1S) and 65.41% (10D10S)—indicating better recovery at higher sucrose the 21–30 µm class (10D10S vs. 10D0S: OR = 2.11, 95% CI = 1.44–3.10; 10D10S vs. 10D1S: OR = 1.82, 95% CI = 1.25–2.64).

### Neurula

For neurulas (Fig. 3C), the 11–20 µm class declined post-thaw for 10D0S (34.41%, OR: 3.12, 95% CI: 1.8–5.41), 10D1S (28.94%, OR: 4.02, 95% CI: 2.33–6.94), and 10D10S (30.19%, OR: 3.79, 95% CI: 2.21–6.49) compared to pre-freeze (62.09%). Conversely, the 21–30 µm class increased for 10D0S (46.07%, OR: 0.51, 95% CI: 0.29–0.88), 10D1S (55.69%, OR: 0.35, 95% CI: 0.20–0.60), and 10D10S (58.38%, OR: 0.31, 95% CI: 0.18–0.53) relative to pre-freeze (30.37%). Recovery was higher in 10D10S than 10D0S (OR: 1.64, 95% CI: 1.15–2.35), but not different from 10D1S (OR: 1.12, 95% CI: 0.79–1.59).

### R. littlejohni neurula cells cryopreserved with 10D10S

Post-thaw cell concentration decreased significantly from 338,422 to 237,403 cells/mL (LRT χ²(1) = 8.97, P = 0.002; OR = 8.55, 95% CI: 6.94–10.54; Fig 7A). Membrane-intact cells also declined significantly from 91.87% to 46.69% (LRT χ²(1) = 110.09, P = 9.34 × 10⁻²□; OR = 12.89, 95% CI: 9.97–16.68; Fig. 7B).

**Figure 6:**
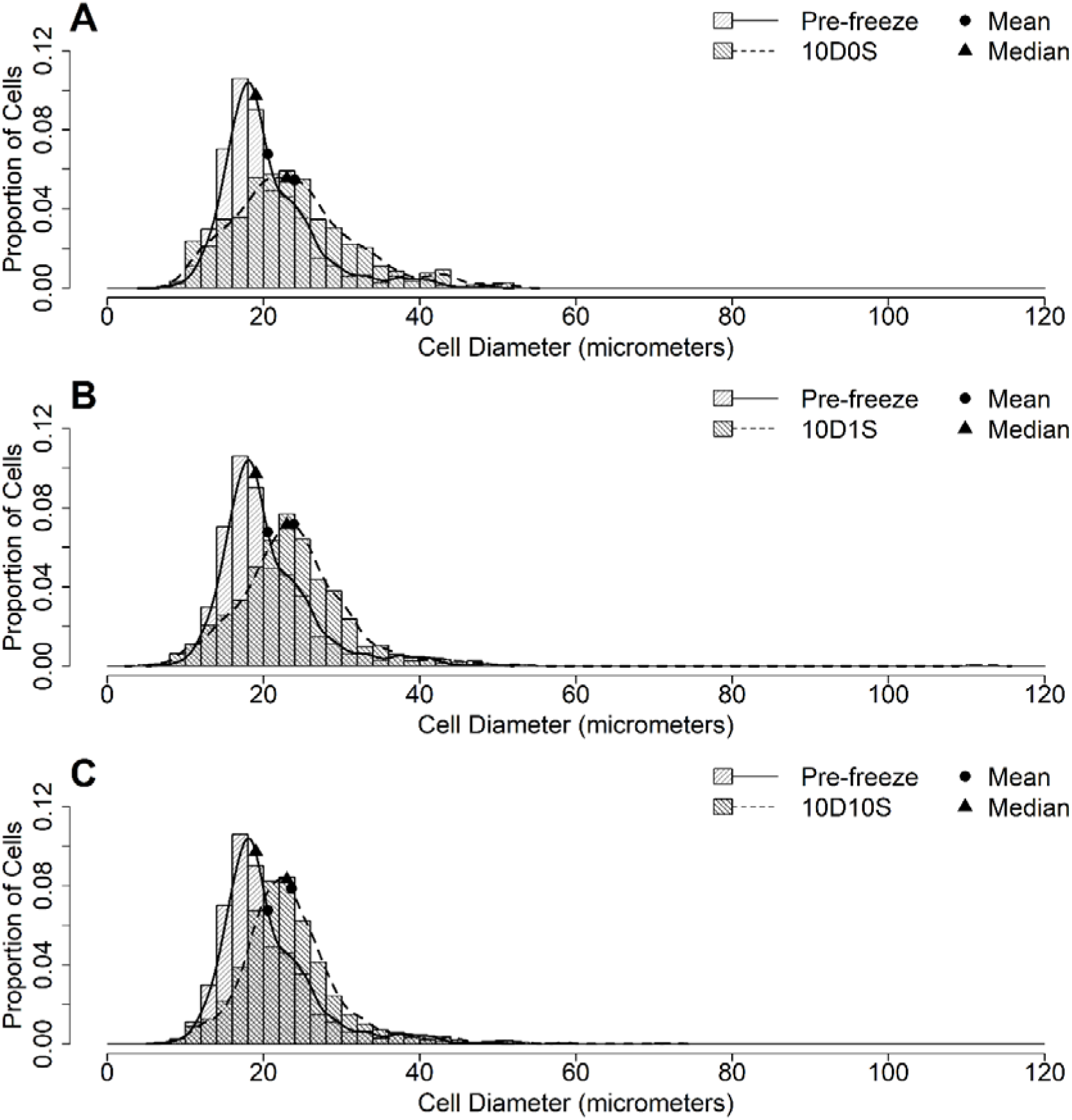
Cell size distributions of *L. peronii* neurulas pre-freeze (mean 20.53 µm, median 19 µm) and post-thaw in (A) 10D0S (mean 23.99 µm), (B) 10D1S (mean 23.89 µm), and (C) 10D10S (mean 23.6 µm).

**Figure 7.**
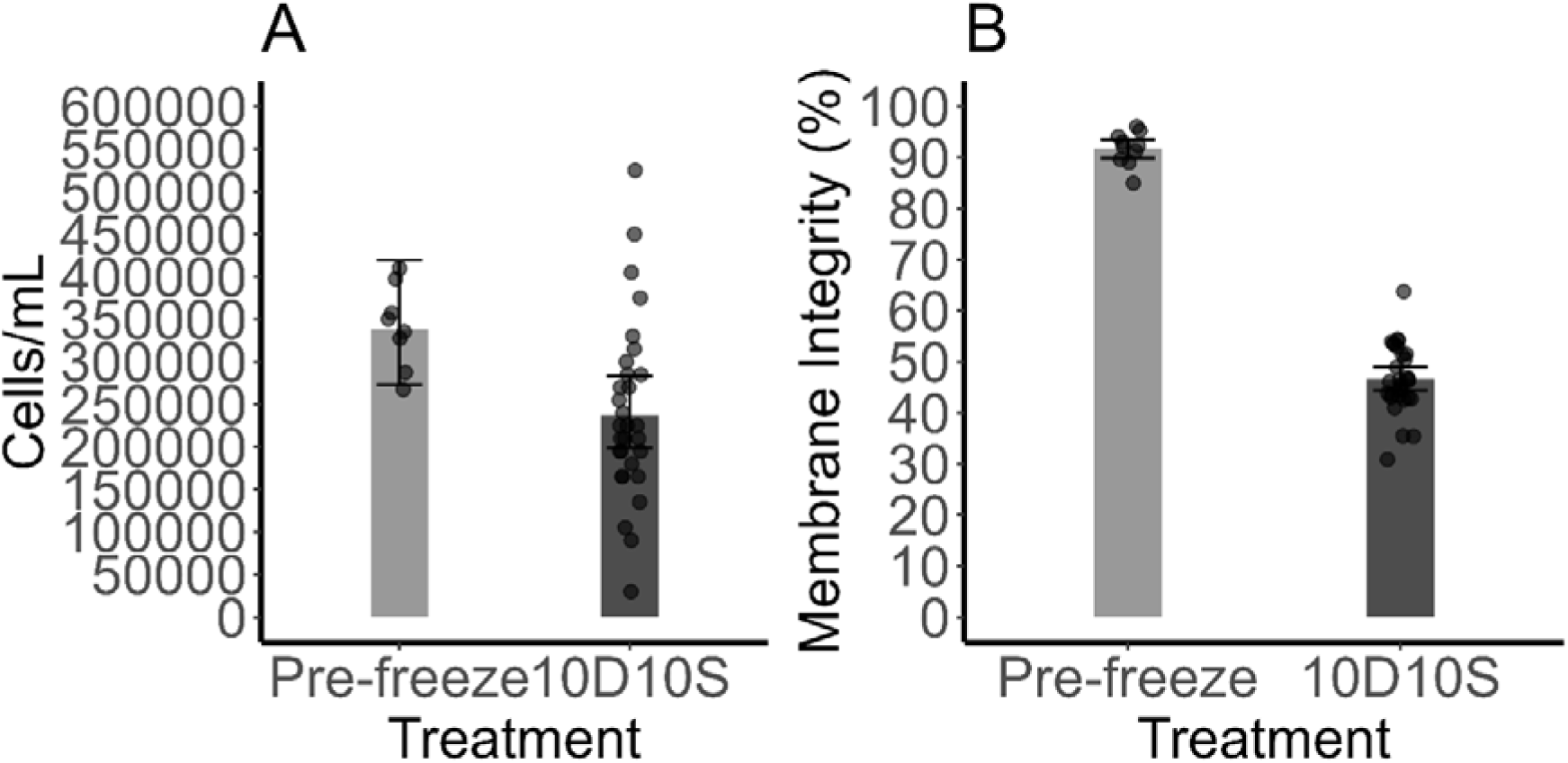
R. littlejohni neurula cells cryopreserved with 10D10S. (A) Cell concentration (binomial GLMM) and (B) % membrane-intact cells pre-freeze vs. post-thaw (negative binomial GLMM). EMMs with 95% CI and raw data (dots) shown.

#### Cell size and cell size class distributions pre-freeze and post-thaw

Post-thaw cell diameters significantly shifted toward larger cells compared to pre-freeze (D = 0.2, P = 3.43×10⁻¹²; Figure 11), similar to *L. peronii* neurulas (Fig. 8). A significant effect of size class × CPA interaction (LRT χ²(8) = 20.41, P = 0.009; Fig. 9) between pre-freeze and 10D10S. There were significantly fewer 11–20 µm cells were recovered post-thaw for 10D10S (19.4%) compared to pre-freeze (34.1%) (OR = 2.16, 95% CI: 1.23–3.79). There was a significant increase in the percentage of cells recovered in the 31-40 µm class post-thaw (14.5%) than pre-freeze (7.5%, OR: 0.48, 95% CI: 0.26-0.89) and in the 41-50 µm cell class post-thaw (4.23%) than pre-freeze (1.43%, OR: 0.33, 95% CI: 0.13-0.82).

**Figure 8:**
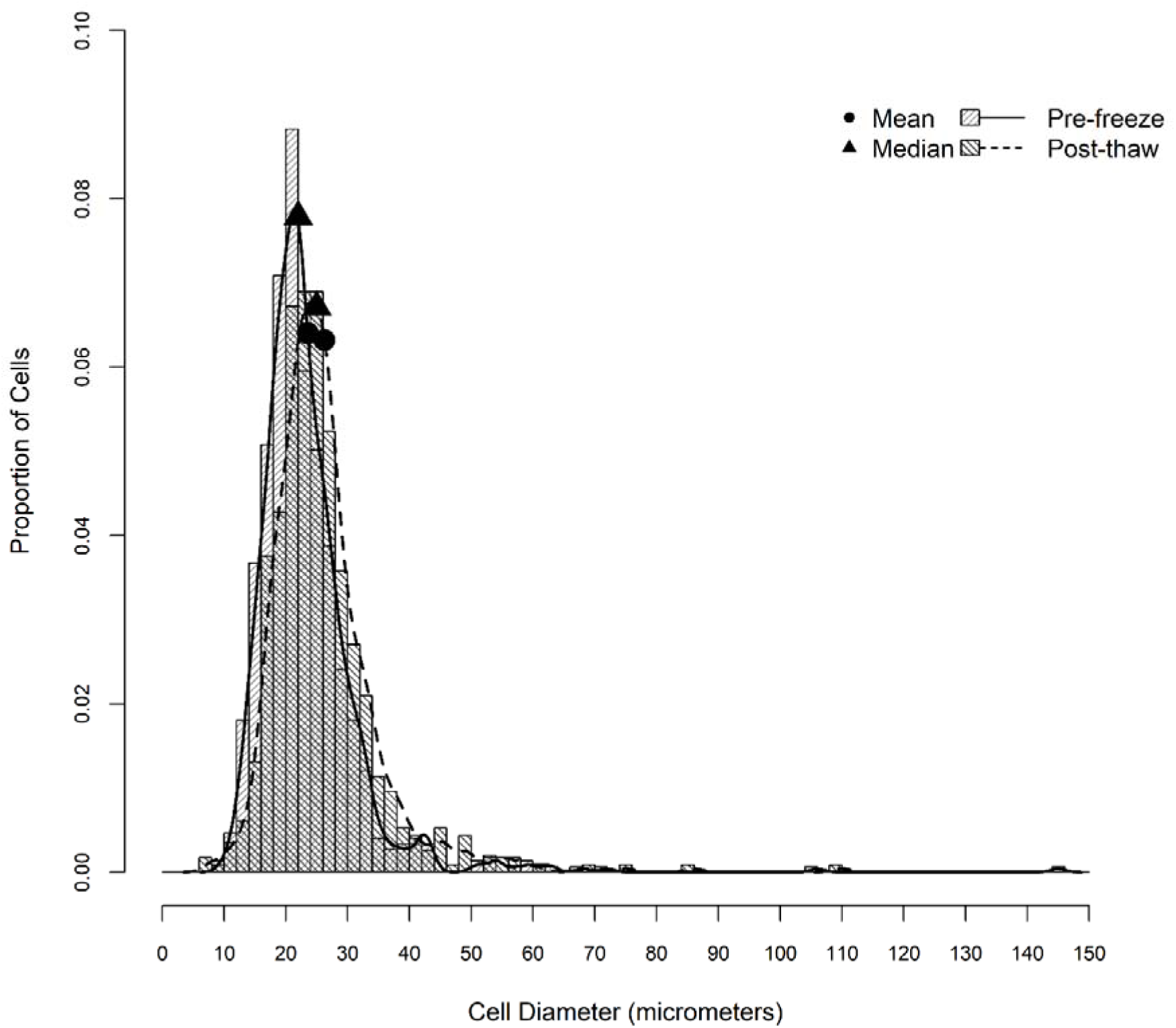
Cell size distributions for *R. littlejohni* neurula cell preparations pre-freeze and post-thaw. Means and medians for cell size are 23.6 µm and 22 µm (pre-freeze), and 26.2 µm and 25 µm (post-thaw), respectively.

**Figure 9:**
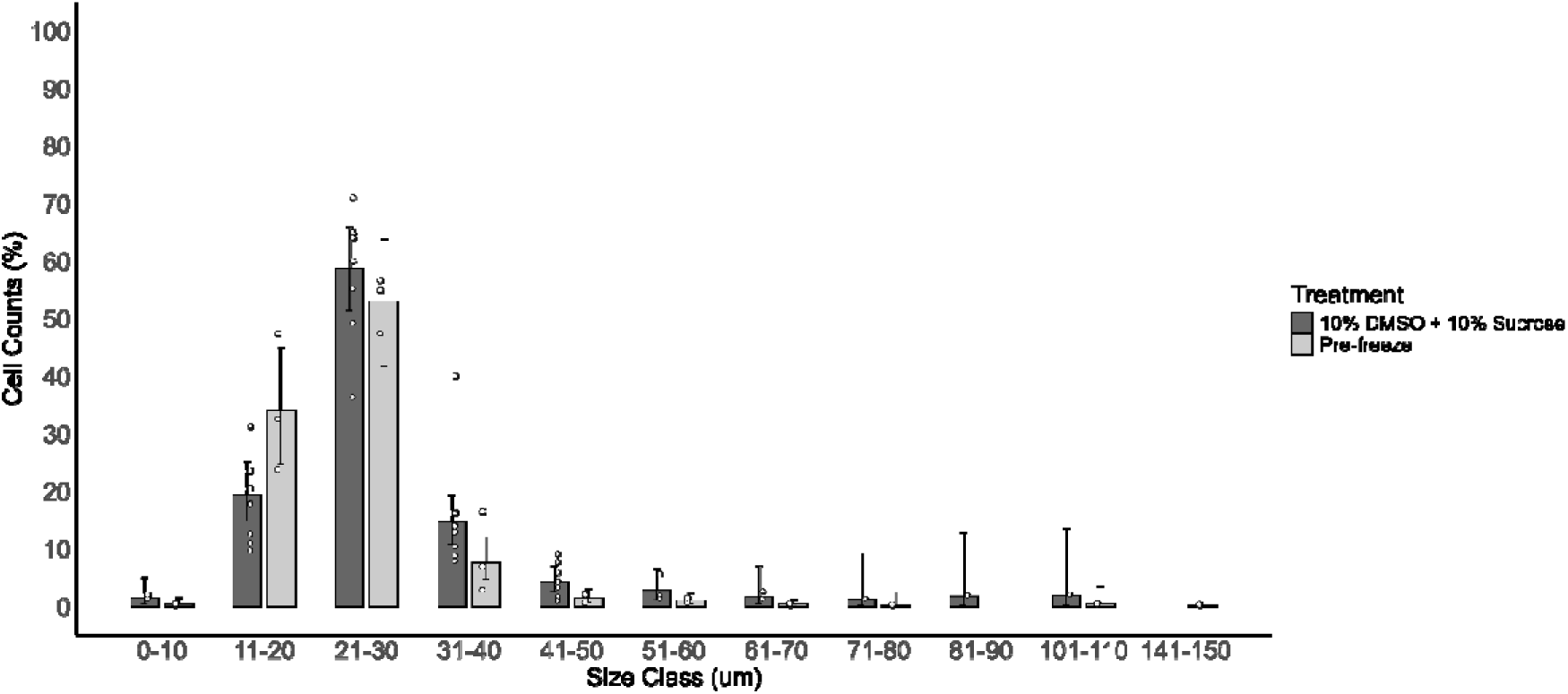
Percentage of cell counts in various size classes pre-freeze and post-thaw for *R. littlejohni*. GLMM binomial. EMMs are plotted along with 95% CIs and raw data (dots).

## Discussion

This study demonstrated for the first time the importance of cell size distribution in the cryopreservation response of embryonic cells in vertebrates with large eggs and embryos. Recovery and viability were quantified across cells of differing sizes under constant cryopreservation conditions, where cryoprotectant composition and cooling rates were held constant, and cell size distribution was treated as the response variable. The differential response of cells to freezing and thawing provides a new explanation for why intact embryos fail to survive slow cooling—termed here the Goldilocks effect, where conditions optimal for one cell size are suboptimal for others.

Cell size distribution is rarely considered in cryopreservation, though it can significantly affect outcomes (Fadda et al., 2010; Fadda et al., 2011; Griffiths et al., 1979). When studied empirically (Köseoğlu et al., 2001; Toner et al., 1992) and modelled in detail (Fadda et al., 2010; Fadda et al., 2011), it is clear that cell size affects important aspects of freezing and thawing through effects including rates of diffusion and fluxes of cryoprotectant and water across the plasmalemma and the temperature at which the cytoplasm freezes (Fadda et al., 2010; Fadda et al., 2011). Cell size is often overlooked in cryopreservation because many populations show little variation (Fadda et al., 2010); however, some cell populations—including rat hepatocytes (25–35 µm)—do vary (Toner et al., 1992).Accounting for cell size distribution may explain much of the variation in cryopreservation responses across cell systems.

In the current study, based on concentration and membrane integrity, we showed that under a constant set of cryopreservation conditions, there was an optimal size for post-thaw recovery and viability. By extension, protocols adjusted to increase recovery of cells at one size will reduce the recovery of other cell sizes. We present a graphical representation of the Goldilocks hypothesis (see Figure 10), which highlights the theoretical basis of the hypothesis and recognises the likely complex effects underpinning it (i.e. that the drivers of decline in recovery above and below the optimum size for a given set of conditions are likely to be different).

**Figure 10.**
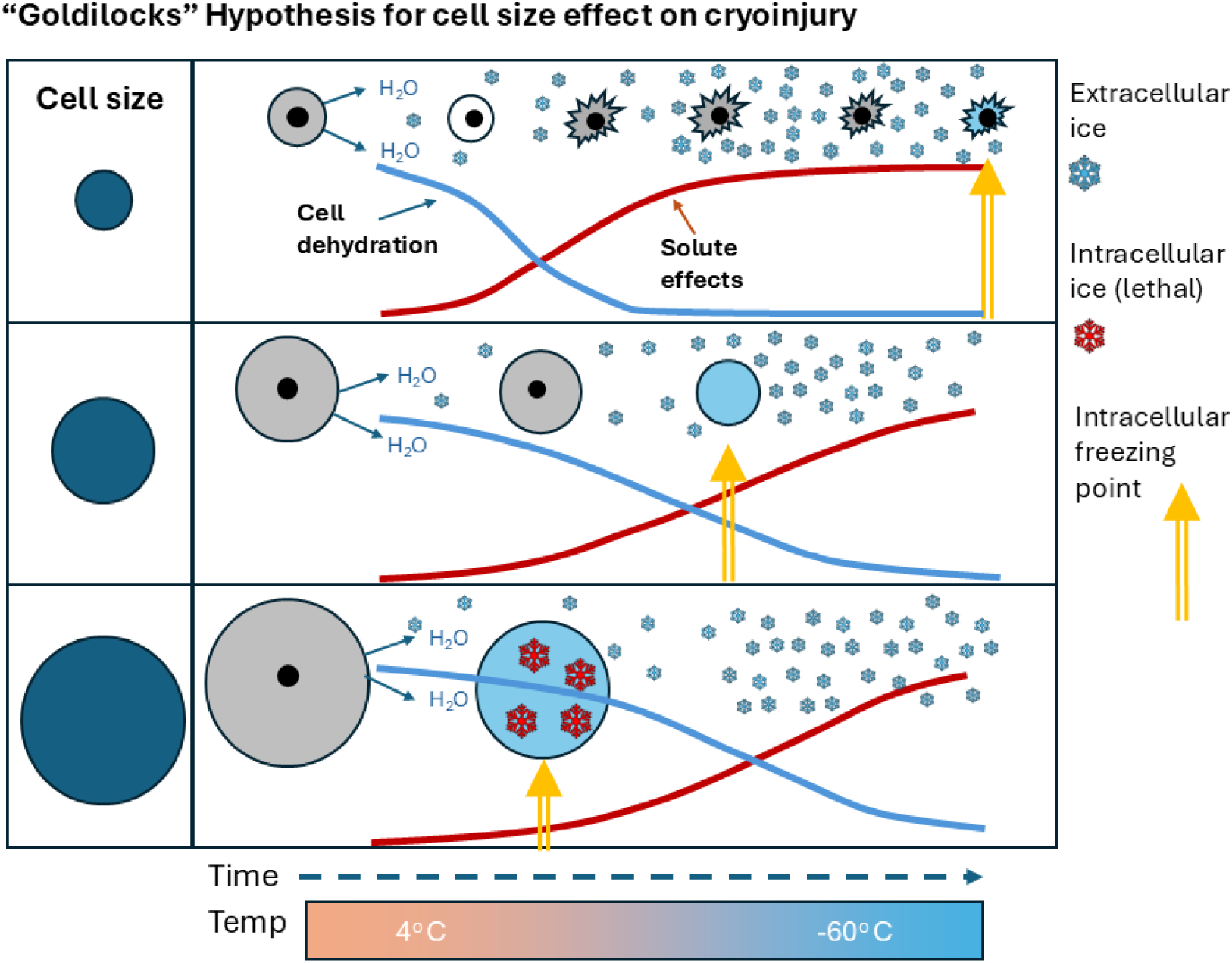
The Goldilocks hypothesis of cryoinjury proposes a differential response of cells to cryopreservation as a function of cell size, with an optimal size yielding the highest survival under given conditions (cryoprotectant, and cooling and thawing rates). Cells smaller than optimal size are prone to excessive dehydration, shrinkage, and osmotic stress, while larger cells experience incomplete dehydration and intracellular ice formation, both leading to reduced recovery.

We suggested that in the case of cells smaller than the optimum size, dehydration proceeds more rapidly due to a larger cell surface area for transport of water and permeating cryoprotectant), but the freezing point is lower (based on theoretical expectations of fewer ice nucleation points in smaller volumes) (Fadda et al., 2010; Fadda et al., 2011). We hypothesise this results in increased cryoinjury from solute effects (Mazur, 1963, 2004) relative to optimal cell size. However, cells larger than the optimal cell size are predicted to dehydrate more slowly (small surface area to volume ratio), but freeze at a higher temperatures (due to the probability of more ice nucleation points in larger cytoplasmic volumes (Fadda et al., 2010; Fadda et al., 2011). Consequently, we hypothesise that lethal intracellular ice formation is more likely the cause of cryoinjury during the cooling of larger cells. The hypothesis derived in this study is essentially another form of the two-factor model of cryoinjury proposed by Peter Mazur (Mazur, 1970; Mazur et al., 1972) in which variations in cooling rates lead to an inverted parabolic pattern of cell recovery. In that model, variation in cooling rate drives the model. In this study, variation in cell size drives a similarly shaped predicted curve.

The current study showed that the recovery of cryopreserved cells was higher in later stages of embryo development (gastrula and neurula compared to blastula), which correlates with an increased frequency of smaller cells in later stages. The most parsimonious explanation for this is that larger cells are more prone to cryoinjury. Similar findings of higher recovery in later developmental stages reported for fish embryos (Calvi & Maisse, 1998; Calvi & Maisse, 1999; Strussmann et al., 1999) were suggested to be either a function of smaller cell size or the slowing cell cycle (fewer dividing cells); however, no direct evidence for either hypothesis was obtained experimentally.

The results with sucrose in this study recognised the best cryoprotectant combination as 10% DMSO and 10% sucrose, which produced the highest recovery of cell concentration (cells/ml) and membrane intact cells across the entire dissociated cell population. Modifications of the protocol would be required to achieve the highest recovery of cells in a different range (e.g. by altering cooling and thawing rates, cryoprotectant selection, and cryoprotectant concentrations).

The addition of sucrose to the cryoprotectant medium improved the post-thaw recovery of cells at all three developmental stages. As the sucrose concentration increased, so did the cell concentration and the percentage of membrane intact cells post-thaw. In other cryopreservation studies, addition of sucrose led to increased recovery post-thaw in zebrafish blastomeres, human oocytes, ram sperm, and human foetal liver cells (Arando et al., 2017; Fabbri et al., 2001; Lin et al., 2009; Petrenko et al., 2008). Sucrose is likely to increase dehydration of cells, which limits IIF. Also, sucrose is known to stabilise membranes, which reduces solute effects during extracellular ice formation (Leekumjorn & Sum, 2008; Tsai et al., 2018). Hence, it is likely to both reduce IIF in larger cells and reduce solute effects in smaller cells, contributing by two separate mechanisms to the Goldilocks effect. In a previous study on *L. peronii* (Lawson et al., 2013), there was low post-thaw membrane integrity of gastrula and neurula cells cryopreserved without sucrose (4.4% with 10% DMSO for gastrula and 5.1% with 15% glycerol for neurula). In comparison, 10D10S achieved 74.074% and 69.490% post-thaw recovery of membrane intact cells in gastrulas and neurulas, respectively.

Even protocols based on vitrification of amphibian (Clulow et al., 2019) and fish embryos (Hagedorn et al., 1997, 1997, & 1998) have failed to prevent IIF during cryopreservation. On theoretical grounds supported by experimental evidence (Khosla et al., 2017; Khosla et al., 2018a; Khosla et al., 2018b), it is thought that most of the intracellular ice crystal formation occurs during warming (thawing) even when vitrification is rapid enough to prevent IIF during freezing. A recent breakthrough utilising super rapid warming employing laser pulses has allowed the recovery of live embryos in the zebrafish (Khosla et al., 2020), which may provide a solution to cryopreservation of small eggs and embryos of some fish and amphibian species. However, the technology is unlikely to be viable (Khosla et al., 2018b) for structures with diameters above a limit of about 2000 µm, suggesting that cryopreservation of dissociated embryos will continue to be the only viable method for anamniotes with eggs and embryos over 2000 µm in diameter (Clulow et al., 2019). In this respect, the current study provides a way forward for the recovery of diploid amphibian genomes by demonstrating that high levels of recovery of slow-cooled cells of particular sizes are possible under conditions where cooling rates and non-permeating cryoprotectant concentrations are optimised. A sub-set of embryonic cells are thus likely to be able to be recovered as viable embryos when combined with either nuclear transfer or the generation of chimeras in amphibians (Clulow & Clulow, 2016; Clulow et al., 2019).

The current study provides a strong evidence base for optimising recovery of diploid genomes of anamniotes generally, but especially amphibians which are so far poorly researched (Lawson et al., 2013). The utility of slow cooling cryopreservation of dissociated embryonic cells has also been demonstrated in various fish species in which recovery of cryopreserved blastomeres have been reported (Calvi & Maisse, 1998; Calvi & Maisse, 1999; Kusuda et al., 2004; Lin et al., 2009; Nilsson & Cloud, 1993; Strussmann et al., 1999) followed by the generation of live offspring from cryopreserved blastomeres through the generation of chimeras by cell transplantation (Calvi & Maisse, 1998; Calvi & Maisse, 1999; Kusuda et al., 2004; Strussmann et al., 1999; Uteshev et al., 2002; Yoshizaki & Lee, 2018).

The generality of these results is supported by the successful recovery of slow-cooled embryonic cells from the threatened *R. littlejohni*. Recovery patterns mirrored those in *L. peronii*, despite phylogenetic distance, suggesting a cell size–based response rather than a phylogenetic effect. This represents the first report of live embryonic cell recovery from a threatened amphibian, indicating that the approach may be valuable as conservation tool for amphibian biobanking.

