## Supplementary files for "Goldilocks conundrum explains cryoinjury in slow-cooled amphibian embryonic cells"

***Supplementary* *Materials***

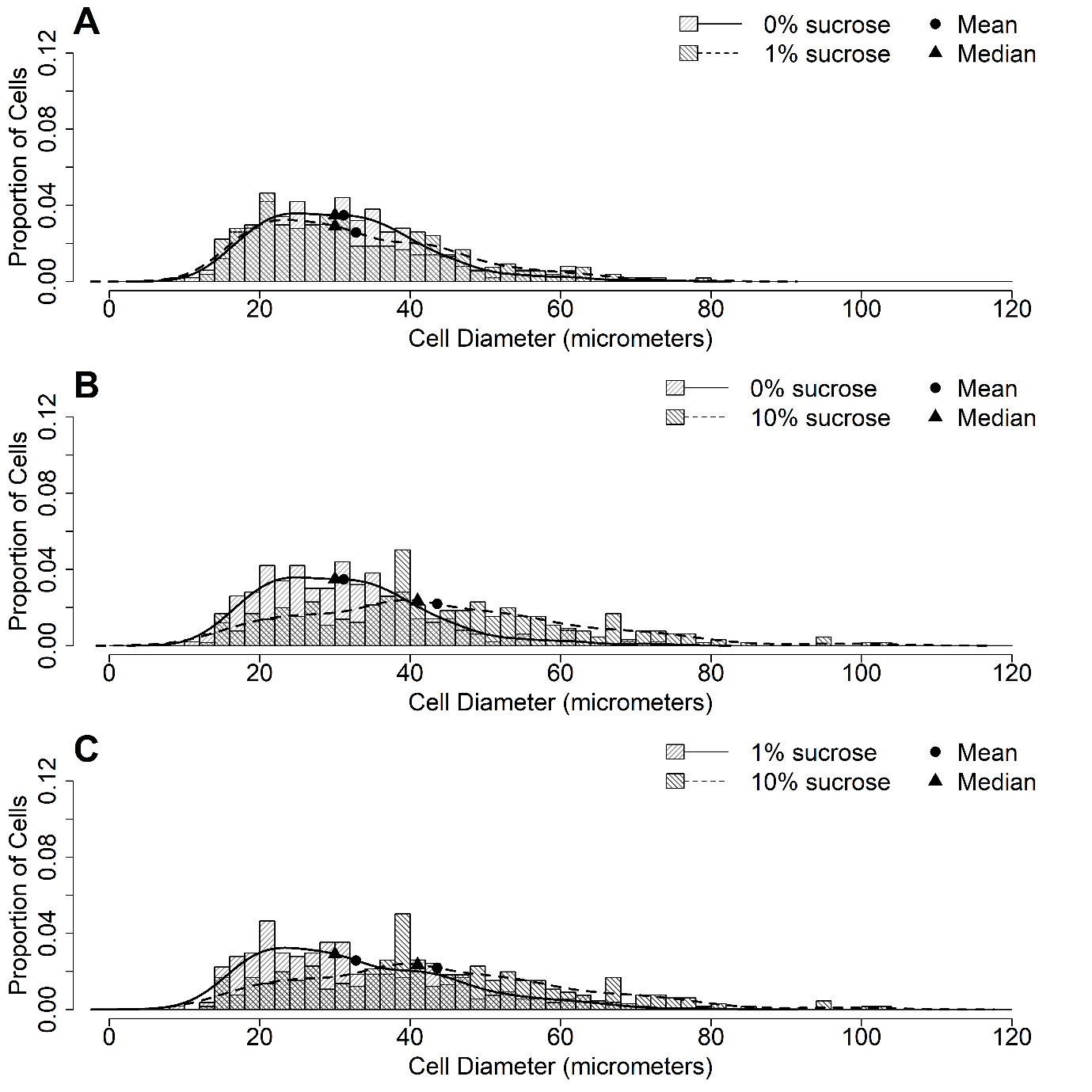

Supplementary figure 1: *L. peronii* blastula post-thaw cell size distributions showing direct comparisons between 0% (mean: 31.2 µm and median: 30 µm) sucrose and 1% sucrose (mean: 32.8 µm and median: 30 µm) (A), 0% sucrose and 10% sucrose (1B), and 1% sucrose to 10% sucrose (mean: 43.6 µm and median: 41 µm) (C).

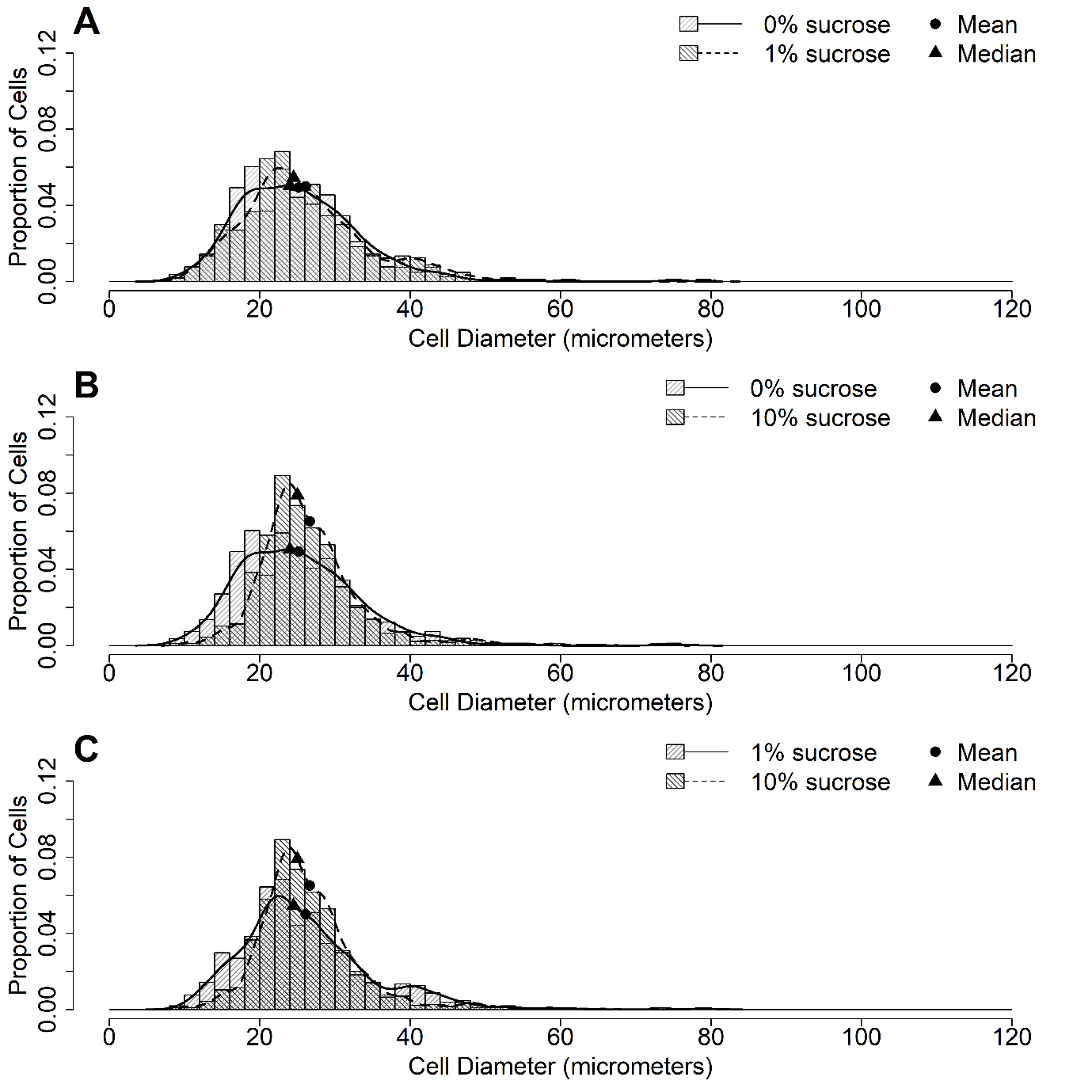

Supplementary figure 2: *L. peronii* gastrula post-thaw cell size distributions showing direct comparisons between 0% sucrose (mean: 25.2 µm and median: 24 µm) and 1% sucrose (mean: 26.1 µm and median: 25 µm) (A), 0% sucrose and 10% sucrose (mean: 26.7 µm and median: 25 µm) (B), and 1% sucrose and 10% sucrose (C).

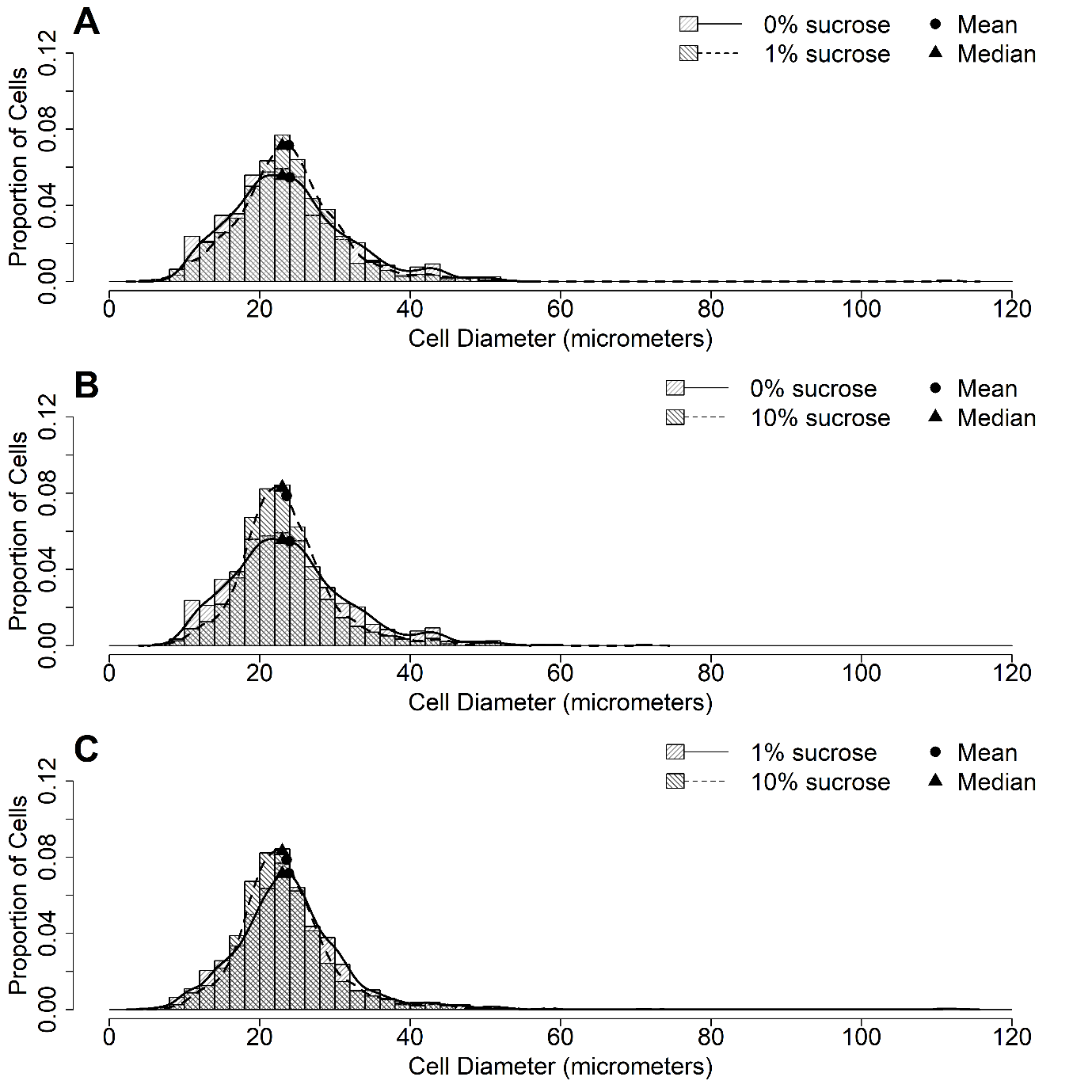
 Supplementary figure 3: *L. peronii* neurula post-thaw cell size distributions showing direct comparisons between 0% and 1% sucrose (mean: 24.0 µm and median: 23 µm) (mean: 23.9 µm and median: 23 µm) (A), 0% and 10% sucrose (mean: 23.6 µm and median: 23 µm) (B), and 1% and 10% sucrose (C).

Supplementary table 1: Odds ratios and 95% confidence intervals for percentage of membrane intact cells for pairwise comparisons between all possible cryoprotectant and stage (blastula, gastrula, and neurula) combinations in *L. peronii.*

| Membrane integrity | | | | | | | |
| --- | --- | --- | --- | --- | --- | --- | --- |
| reverse contrast | odds ratio | LCL^1^ | UCL^2^ | contrast | odds ratio | LCL^1^ | UCL^2^ |
| 10D1S blast / 10D0S blast | 1.349* | 1.136 | 1.604 | 10D0S blast / 10D1S blast | 0.741* | 0.624 | 0.881 |
| 10D10S blast / 10D0S blast | 2.223* | 1.871 | 2.641 | 10D0S blast / 10D10S blast | 0.45* | 0.379 | 0.534 |
| 10D10S blast / 10D1S blast | 1.647* | 1.387 | 1.957 | 10D0S blast / pre-freeze blast | 0.042* | 0.03 | 0.059 |
| pre-freeze blast / 10D0S blast | 23.733* | 17.012 | 33.109 | 10D0S blast / 10D0S gast | 0.272* | 0.227 | 0.327 |
| pre-freeze blast / 10D1S blast | 17.588* | 12.606 | 24.537 | 10D0S blast / 10D1S gast | 0.202* | 0.157 | 0.259 |
| pre-freeze blast / 10D10S blast | 10.676* | 7.658 | 14.882 | 10D0S blast / 10D10S gast | 0.122* | 0.095 | 0.158 |
| 10D0S gast / 10D0S blast | 3.675* | 3.062 | 4.411 | 10D0S blast / pre-freeze gast | 0.011* | 0.008 | 0.017 |
| 10D0S gast / 10D1S blast | 2.723* | 2.117 | 3.503 | 10D0S blast / 10D0S neur | 0.341* | 0.285 | 0.409 |
| 10D0S gast / 10D10S blast | 1.653* | 1.291 | 2.117 | 10D0S blast / 10D1S neur | 0.253* | 0.197 | 0.324 |
| 10D0S gast / pre-freeze blast | 0.155* | 0.107 | 0.223 | 10D0S blast / 10D10S neur | 0.154* | 0.119 | 0.197 |
| 10D1S gast / 10D0S blast | 4.959* | 3.86 | 6.372 | 10D0S blast / pre-freeze neur | 0.014* | 0.01 | 0.021 |
| 10D1S gast / 10D1S blast | 3.675* | 3.062 | 4.411 | 10D1S blast / 10D10S blast | 0.607* | 0.511 | 0.721 |
| 10D1S gast / 10D10S blast | 2.231* | 1.743 | 2.856 | 10D1S blast / pre-freeze blast | 0.057* | 0.041 | 0.079 |
| 10D1S gast / pre-freeze blast | 0.209* | 0.145 | 0.301 | 10D1S blast / 10D0S gast | 0.367* | 0.285 | 0.472 |
| 10D1S gast / 10D0S gast | 1.349* | 1.136 | 1.604 | 10D1S blast / 10D1S gast | 0.272* | 0.227 | 0.327 |
| 10D10S gast / 10D0S blast | 8.17* | 6.333 | 10.539 | 10D1S blast / 10D10S gast | 0.165* | 0.128 | 0.213 |
| 10D10S gast / 10D1S blast | 6.054* | 4.691 | 7.815 | 10D1S blast / pre-freeze gast | 0.015* | 0.01 | 0.023 |
| 10D10S gast / 10D10S blast | 3.675* | 3.062 | 4.411 | 10D1S blast / 10D0S neur | 0.461* | 0.359 | 0.591 |
| 10D10S gast / pre-freeze blast | 0.344* | 0.238 | 0.497 | 10D1S blast / 10D1S neur | 0.341* | 0.285 | 0.409 |
| 10D10S gast / 10D0S gast | 2.223* | 1.871 | 2.641 | 10D1S blast / 10D10S neur | 0.207* | 0.161 | 0.267 |
| 10D10S gast / 10D1S gast | 1.647* | 1.387 | 1.957 | 10D1S blast / pre-freeze neur | 0.019* | 0.013 | 0.029 |
| pre-freeze gast / 10D0S blast | 87.22* | 58.907 | 129.142 | 10D10S blast / pre-freeze blast | 0.094* | 0.067 | 0.131 |
| pre-freeze gast / 10D1S blast | 64.636* | 43.635 | 95.743 | 10D10S blast / 10D0S gast | 0.605* | 0.472 | 0.775 |
| pre-freeze gast / 10D10S blast | 39.234* | 26.577 | 57.92 | 10D10S blast / 10D1S gast | 0.448* | 0.35 | 0.574 |
| pre-freeze gast / pre-freeze blast | 3.675* | 3.062 | 4.411 | 10D10S blast / 10D10S gast | 0.272* | 0.227 | 0.327 |
| pre-freeze gast / 10D0S gast | 23.733* | 17.012 | 33.109 | 10D10S blast / pre-freeze gast | 0.025* | 0.017 | 0.038 |
| pre-freeze gast / 10D1S gast | 17.588* | 12.606 | 24.537 | 10D10S blast / 10D0S neur | 0.759* | 0.593 | 0.971 |
| pre-freeze gast / 10D10S gast | 10.676* | 7.658 | 14.882 | 10D10S blast / 10D1S neur | 0.562* | 0.44 | 0.719 |
| 10D0S neur / 10D0S blast | 2.93* | 2.448 | 3.506 | 10D10S blast / 10D10S neur | 0.341* | 0.285 | 0.409 |
| 10D0S neur / 10D1S blast | 2.171* | 1.691 | 2.787 | 10D10S blast / pre-freeze neur | 0.032* | 0.022 | 0.047 |
| 10D0S neur / 10D10S blast | 1.318* | 1.03 | 1.686 | pre-freeze blast / 10D0S gast | 6.458* | 4.476 | 9.316 |
| 10D0S neur / pre-freeze blast | 0.123* | 0.085 | 0.178 | pre-freeze blast / 10D1S gast | 4.786* | 3.318 | 6.902 |
| 10D0S neur / 10D0S gast | 0.797* | 0.681 | 0.933 | pre-freeze blast / 10D10S gast | 2.905* | 2.01 | 4.198 |
| 10D0S neur / 10D1S gast | 0.591* | 0.468 | 0.746 | pre-freeze blast / pre-freeze gast | 0.272* | 0.227 | 0.327 |
| 10D0S neur / 10D10S gast | 0.359* | 0.284 | 0.453 | pre-freeze blast / 10D0S neur | 8.101* | 5.606 | 11.705 |
| 10D0S neur / pre-freeze gast | 0.034* | 0.023 | 0.049 | pre-freeze blast / 10D1S neur | 6.003* | 4.156 | 8.672 |
| 10D1S neur / 10D0S blast | 3.953* | 3.083 | 5.069 | pre-freeze blast / 10D10S neur | 3.644* | 2.519 | 5.271 |
| 10D1S neur / 10D1S blast | 2.93* | 2.448 | 3.506 | pre-freeze blast / pre-freeze neur | 0.341* | 0.285 | 0.409 |
| 10D1S neur / 10D10S blast | 1.778* | 1.391 | 2.274 | 10D0S gast / 10D1S gast | 0.741* | 0.624 | 0.881 |
| 10D1S neur / pre-freeze blast | 0.167* | 0.115 | 0.241 | 10D0S gast / 10D10S gast | 0.45* | 0.379 | 0.534 |
| 10D1S neur / 10D0S gast | 1.076 | 0.852 | 1.359 | 10D0S gast / pre-freeze gast | 0.042* | 0.03 | 0.059 |
| 10D1S neur / 10D1S gast | 0.797* | 0.681 | 0.933 | 10D0S gast / 10D0S neur | 1.254* | 1.072 | 1.468 |
| 10D1S neur / 10D10S gast | 0.484* | 0.383 | 0.612 | 10D0S gast / 10D1S neur | 0.93 | 0.736 | 1.174 |
| 10D1S neur / pre-freeze gast | 0.045* | 0.031 | 0.066 | 10D0S gast / 10D10S neur | 0.564* | 0.447 | 0.712 |
| 10D1S neur / 10D0S neur | 1.349* | 1.136 | 1.604 | 10D0S gast / pre-freeze neur | 0.053* | 0.037 | 0.076 |
| 10D10S neur / 10D0S blast | 6.513* | 5.065 | 8.375 | 10D1S gast / 10D10S gast | 0.607* | 0.511 | 0.721 |
| 10D10S neur / 10D1S blast | 4.826* | 3.751 | 6.21 | 10D1S gast / pre-freeze gast | 0.057* | 0.041 | 0.079 |
| 10D10S neur / 10D10S blast | 2.93* | 2.448 | 3.506 | 10D1S gast / 10D0S neur | 1.693* | 1.34 | 2.138 |
| 10D10S neur / pre-freeze blast | 0.274* | 0.19 | 0.397 | 10D1S gast / 10D1S neur | 1.254* | 1.072 | 1.468 |
| 10D10S neur / 10D0S gast | 1.772* | 1.405 | 2.236 | 10D1S gast / 10D10S neur | 0.761* | 0.604 | 0.961 |
| 10D10S neur / 10D1S gast | 1.313* | 1.041 | 1.657 | 10D1S gast / pre-freeze neur | 0.071* | 0.049 | 0.103 |
| 10D10S neur / 10D10S gast | 0.797* | 0.681 | 0.933 | 10D10S gast / pre-freeze gast | 0.094* | 0.067 | 0.131 |
| 10D10S neur / pre-freeze gast | 0.075* | 0.052 | 0.108 | 10D10S gast / 10D0S neur | 2.789* | 2.206 | 3.526 |
| 10D10S neur / 10D0S neur | 2.223* | 1.871 | 2.641 | 10D10S gast / 10D1S neur | 2.067* | 1.634 | 2.613 |
| 10D10S neur / 10D1S neur | 1.647* | 1.387 | 1.957 | 10D10S gast / 10D10S neur | 1.254* | 1.072 | 1.468 |
| pre-freeze neur / 10D0S blast | 69.53* | 47.157 | 102.519 | 10D10S gast / pre-freeze neur | 0.118* | 0.082 | 0.169 |
| pre-freeze neur / 10D1S blast | 51.526* | 34.932 | 76.003 | pre-freeze gast / 10D0S neur | 29.771* | 20.537 | 43.157 |
| pre-freeze neur / 10D10S blast | 31.277* | 21.261 | 46.011 | pre-freeze gast / 10D1S neur | 22.062* | 15.218 | 31.985 |
| pre-freeze neur / pre-freeze blast | 2.93* | 2.448 | 3.506 | pre-freeze gast / 10D10S neur | 13.392* | 9.251 | 19.386 |
| pre-freeze neur / 10D0S gast | 18.919* | 13.131 | 27.26 | pre-freeze gast / pre-freeze neur | 1.254* | 1.072 | 1.468 |
| pre-freeze neur / 10D1S gast | 14.02* | 9.73 | 20.202 | 10D0S neur / 10D1S neur | 0.741* | 0.624 | 0.881 |
| pre-freeze neur / 10D10S gast | 8.511* | 5.906 | 12.263 | 10D0S neur / 10D10S neur | 0.45* | 0.379 | 0.534 |
| pre-freeze neur / pre-freeze gast | 0.797* | 0.681 | 0.933 | 10D0S neur / pre-freeze neur | 0.042* | 0.03 | 0.059 |
| pre-freeze neur / 10D0S neur | 23.733* | 17.012 | 33.109 | 10D1S neur / 10D10S neur | 0.607* | 0.511 | 0.721 |
| pre-freeze neur / 10D1S neur | 17.588* | 12.606 | 24.537 | 10D1S neur / pre-freeze neur | 0.057* | 0.041 | 0.079 |
| pre-freeze neur / 10D10S neur | 10.676* | 7.658 | 14.882 | 10D10S neur / pre-freeze neur | 0.094* | 0.067 | 0.131 |

^1^Lower confidence limit, ^2^Upper confidence limit, * Significant pairwise comparisons

Supplementary Table 2: Odds ratios and 95% confidence intervals for cell concentration for pairwise comparisons between all possible cryoprotectant and stage (blastula, gastrula, and neurula) combinations for *L. peronii.*

| Concentration (cells/mL) | | | | | | | |
| --- | --- | --- | --- | --- | --- | --- | --- |
| reverse contrast | Odds ratio | LCL^1^ | UCL^2^ | contrast | Odds ratio | LCL^1^ | UCL^2^ |
| 10D1S blast / 10D0S blast | 1.246* | 1.070 | 1.450 | 10D0S blast / 10D1S blast | 0.803* | 0.690 | 0.934 |
| 10D10S blast / 10D0S blast | 1.880* | 1.623 | 2.178 | 10D0S blast / 10D10S blast | 0.532* | 0.459 | 0.616 |
| 10D10S blast / 10D1S blast | 1.509* | 1.308 | 1.741 | 10D0S blast / pre-freeze blast | 0.061* | 0.049 | 0.076 |
| pre-freeze blast / 10D0S blast | 16.360* | 13.144 | 20.364 | 10D0S blast / 10D0S gast | 0.194* | 0.166 | 0.228 |
| pre-freeze blast / 10D1S blast | 13.133* | 10.578 | 16.306 | 10D0S blast / 10D1S gast | 0.156* | 0.125 | 0.195 |
| pre-freeze blast / 10D10S blast | 8.703* | 7.033 | 10.770 | 10D0S blast / 10D10S gast | 0.103* | 0.083 | 0.129 |
| 10D0S gast / 10D0S blast | 5.143* | 4.386 | 6.031 | 10D0S blast / pre-freeze gast | 0.012* | 0.009 | 0.016 |
| 10D0S gast / 10D1S blast | 4.129* | 3.323 | 5.130 | 10D0S blast / 10D0S neur | 0.114* | 0.097 | 0.133 |
| 10D0S gast / 10D10S blast | 2.736* | 2.206 | 3.393 | 10D0S blast / 10D1S neur | 0.091* | 0.073 | 0.113 |
| 10D0S gast / pre-freeze blast | 0.314* | 0.241 | 0.410 | 10D0S blast / 10D10S neur | 0.061* | 0.049 | 0.075 |
| 10D1S gast / 10D0S blast | 6.407* | 5.128 | 8.004 | 10D0S blast / pre-freeze neur | 0.007* | 0.005 | 0.009 |
| 10D1S gast / 10D1S blast | 5.143* | 4.386 | 6.031 | 10D1S blast / 10D10S blast | 0.663* | 0.574 | 0.764 |
| 10D1S gast / 10D10S blast | 3.408* | 2.748 | 4.227 | 10D1S blast / pre-freeze blast | 0.076* | 0.061 | 0.095 |
| 10D1S gast / pre-freeze blast | 0.392* | 0.300 | 0.511 | 10D1S blast / 10D0S gast | 0.242* | 0.195 | 0.301 |
| 10D1S gast / 10D0S gast | 1.246* | 1.070 | 1.450 | 10D1S blast / 10D1S gast | 0.194* | 0.166 | 0.228 |
| 10D10S gast / 10D0S blast | 9.668* | 7.772 | 12.025 | 10D1S blast / 10D10S gast | 0.129* | 0.104 | 0.159 |
| 10D10S gast / 10D1S blast | 7.761* | 6.275 | 9.599 | 10D1S blast / pre-freeze gast | 0.015* | 0.011 | 0.019 |
| 10D10S gast / 10D10S blast | 5.143* | 4.386 | 6.031 | 10D1S blast / 10D0S neur | 0.142* | 0.114 | 0.176 |
| 10D10S gast / pre-freeze blast | 0.591* | 0.455 | 0.767 | 10D1S blast / 10D1S neur | 0.114* | 0.097 | 0.133 |
| 10D10S gast / 10D0S gast | 1.880* | 1.623 | 2.178 | 10D1S blast / 10D10S neur | 0.075* | 0.061 | 0.093 |
| 10D10S gast / 10D1S gast | 1.509* | 1.308 | 1.741 | 10D1S blast / pre-freeze neur | 0.009* | 0.007 | 0.011 |
| pre-freeze gast / 10D0S blast | 84.140* | 63.827 | 110.916 | 10D10S blast / pre-freeze blast | 0.115* | 0.093 | 0.142 |
| pre-freeze gast / 10D1S blast | 67.544* | 51.454 | 88.667 | 10D10S blast / 10D0S gast | 0.366* | 0.295 | 0.453 |
| pre-freeze gast / 10D10S blast | 44.759* | 34.149 | 58.666 | 10D10S blast / 10D1S gast | 0.293* | 0.237 | 0.364 |
| pre-freeze gast / pre-freeze blast | 5.143* | 4.386 | 6.031 | 10D10S blast / 10D10S gast | 0.194* | 0.166 | 0.228 |
| pre-freeze gast / 10D0S gast | 16.360* | 13.144 | 20.364 | 10D10S blast / pre-freeze gast | 0.022* | 0.017 | 0.029 |
| pre-freeze gast / 10D1S gast | 13.133* | 10.578 | 16.306 | 10D10S blast / 10D0S neur | 0.214* | 0.172 | 0.265 |
| pre-freeze gast / 10D10S gast | 8.703* | 7.033 | 10.770 | 10D10S blast / 10D1S neur | 0.172* | 0.139 | 0.212 |
| 10D0S neur / 10D0S blast | 8.792* | 7.512 | 10.289 | 10D10S blast / 10D10S neur | 0.114* | 0.097 | 0.133 |
| 10D0S neur / 10D1S blast | 7.058* | 5.667 | 8.791 | 10D10S blast / pre-freeze neur | 0.013* | 0.010 | 0.017 |
| 10D0S neur / 10D10S blast | 4.677* | 3.767 | 5.806 | pre-freeze blast / 10D0S gast | 3.181* | 2.440 | 4.146 |
| 10D0S neur / pre-freeze blast | 0.537* | 0.412 | 0.700 | pre-freeze blast / 10D1S gast | 2.554* | 1.959 | 3.329 |
| 10D0S neur / 10D0S gast | 1.709* | 1.504 | 1.943 | pre-freeze blast / 10D10S gast | 1.692* | 1.303 | 2.198 |
| 10D0S neur / 10D1S gast | 1.372* | 1.121 | 1.680 | pre-freeze blast / pre-freeze gast | 0.194* | 0.166 | 0.228 |
| 10D0S neur / 10D10S gast | 0.909 | 0.746 | 1.108 | pre-freeze blast / 10D0S neur | 1.861* | 1.428 | 2.425 |
| 10D0S neur / pre-freeze gast | 0.104* | 0.081 | 0.135 | pre-freeze blast / 10D1S neur | 1.494* | 1.150 | 1.941 |
| 10D1S neur / 10D0S blast | 10.952* | 8.813 | 13.611 | pre-freeze blast / 10D10S neur | 0.990 | 0.764 | 1.283 |
| 10D1S neur / 10D1S blast | 8.792* | 7.512 | 10.289 | pre-freeze blast / pre-freeze neur | 0.114* | 0.097 | 0.133 |
| 10D1S neur / 10D10S blast | 5.826* | 4.712 | 7.204 | 10D0S gast / 10D1S gast | 0.803* | 0.690 | 0.934 |
| 10D1S neur / pre-freeze blast | 0.669* | 0.515 | 0.870 | 10D0S gast / 10D10S gast | 0.532* | 0.459 | 0.616 |
| 10D1S neur / 10D0S gast | 2.130* | 1.754 | 2.586 | 10D0S gast / pre-freeze gast | 0.061* | 0.049 | 0.076 |
| 10D1S neur / 10D1S gast | 1.709* | 1.504 | 1.943 | 10D0S gast / 10D0S neur | 0.585* | 0.515 | 0.665 |
| 10D1S neur / 10D10S gast | 1.133 | 0.937 | 1.370 | 10D0S gast / 10D1S neur | 0.470* | 0.387 | 0.570 |
| 10D1S neur / pre-freeze gast | 0.130* | 0.101 | 0.167 | 10D0S gast / 10D10S neur | 0.311* | 0.257 | 0.377 |
| 10D1S neur / 10D0S neur | 1.246* | 1.070 | 1.450 | 10D0S gast / pre-freeze neur | 0.036* | 0.028 | 0.046 |
| 10D10S neur / 10D0S blast | 16.527* | 13.338 | 20.479 | 10D1S gast / 10D10S gast | 0.663* | 0.574 | 0.764 |
| 10D10S neur / 10D1S blast | 13.267* | 10.726 | 16.410 | 10D1S gast / pre-freeze gast | 0.076* | 0.061 | 0.095 |
| 10D10S neur / 10D10S blast | 8.792* | 7.512 | 10.289 | 10D1S gast / 10D0S neur | 0.729* | 0.595 | 0.892 |
| 10D10S neur / pre-freeze blast | 1.010 | 0.780 | 1.309 | 10D1S gast / 10D1S neur | 0.585* | 0.515 | 0.665 |
| 10D10S neur / 10D0S gast | 3.214* | 2.652 | 3.895 | 10D1S gast / 10D10S neur | 0.388* | 0.319 | 0.470 |
| 10D10S neur / 10D1S gast | 2.580* | 2.126 | 3.130 | 10D1S gast / pre-freeze neur | 0.045* | 0.035 | 0.057 |
| 10D10S neur / 10D10S gast | 1.709* | 1.504 | 1.943 | 10D10S gast / pre-freeze gast | 0.115* | 0.093 | 0.142 |
| 10D10S neur / pre-freeze gast | 0.196* | 0.153 | 0.252 | 10D10S gast / 10D0S neur | 1.100 | 0.903 | 1.340 |
| 10D10S neur / 10D0S neur | 1.880* | 1.623 | 2.178 | 10D10S gast / 10D1S neur | 0.883 | 0.730 | 1.068 |
| 10D10S neur / 10D1S neur | 1.509* | 1.308 | 1.741 | 10D10S gast / 10D10S neur | 0.585* | 0.515 | 0.665 |
| pre-freeze neur / 10D0S blast | 143.836* | 109.319 | 189.251 | 10D10S gast / pre-freeze neur | 0.067* | 0.052 | 0.086 |
| pre-freeze neur / 10D1S blast | 115.467* | 87.858 | 151.751 | pre-freeze gast / 10D0S neur | 9.570* | 7.421 | 12.342 |
| pre-freeze neur / 10D10S blast | 76.515* | 58.379 | 100.287 | pre-freeze gast / 10D1S neur | 7.683* | 5.990 | 9.853 |
| pre-freeze neur / pre-freeze blast | 8.792* | 7.512 | 10.289 | pre-freeze gast / 10D10S neur | 5.091* | 3.976 | 6.519 |
| pre-freeze neur / 10D0S gast | 27.968* | 21.722 | 36.008 | pre-freeze gast / pre-freeze neur | 0.585* | 0.515 | 0.665 |
| pre-freeze neur / 10D1S gast | 22.451* | 17.418 | 28.939 | 10D0S neur / 10D1S neur | 0.803* | 0.690 | 0.934 |
| pre-freeze neur / 10D10S gast | 14.878* | 11.590 | 19.098 | 10D0S neur / 10D10S neur | 0.532* | 0.459 | 0.616 |
| pre-freeze neur / pre-freeze gast | 1.709* | 1.504 | 1.943 | 10D0S neur / pre-freeze neur | 0.061* | 0.049 | 0.076 |
| pre-freeze neur / 10D0S neur | 16.360* | 13.144 | 20.364 | 10D1S neur / 10D10S neur | 0.663* | 0.574 | 0.764 |
| pre-freeze neur / 10D1S neur | 13.133* | 10.578 | 16.306 | 10D1S neur / pre-freeze neur | 0.076* | 0.061 | 0.095 |
| pre-freeze neur / 10D10S neur | 8.703* | 7.033 | 10.770 | 10D10S neur / pre-freeze neur | 0.115* | 0.093 | 0.142 |

^1^Lower confidence limit, ^2^Upper confidence limit, * Significant pairwise comparisons

Supplementary Table 3: Kolmogorov-Smirnov (KS) tests comparing *L. peronii* pre-freeze cell diameter distributions to the post-thaw cell size distributions for the three cryoprotectants within each of the developmental stages.

| Test | D-Statistic | P-Value |
| --- | --- | --- |
| blast_pre_vs_blast_0 | 0.615 | 2.512X10^-38^ |
| blast_pre_vs_blast_1 | 0.538 | 2.435X10^-30^ |
| blast_pre_vs_blast_10 | 0.234 | 1.355X10^-06^ |
| blast_0_vs_blast_1 | 0.115 | 0.065 |
| blast_0_vs_blast_10 | 0.388 | 6.065X10^-19^ |
| blast_1_vs_blast_10 | 0.315 | 3.479X10^-13^ |
| gast_pre_vs_gast_0 | 0.140 | 1.108X10^-04^ |
| gast_pre_vs_gast_1 | 0.163 | 4.901X10^-07^ |
| gast_pre_vs_gast_10 | 0.238 | 4.657X10^-19^ |
| gast_0_vs_gast_1 | 0.088 | 0.058 |
| gast_0_vs_gast_10 | 0.190 | 3.156X10^-09^ |
| gast_1_vs_gast_10 | 0.136 | 9.069X10^-06^ |
| neur_pre_vs_neur_0 | 0.272 | 6.492X10^-25^ |
| neur_pre_vs_neur_1 | 0.323 | 2.147X10^-41^ |
| neur_pre_vs_neur_10 | 0.316 | 2.101X10^-49^ |
| neur_0_vs_neur_10 | 0.084 | 0.007 |
| neur_1_vs_neur_10 | 0.060 | 0.066 |
| neur_0_vs_neur_1 | 0.054 | 0.284 |

Supplementary Table 4: Estimated Marginal Means (EMMs) of % cell counts in various size classes within cryoprotectant combinations for each stage (blastula, gastrula, and neurula) in *L. peronii.*

| EMMs for cell size class | | | |
| --- | --- | --- | --- |
| Cryoprotectant | stage | size class | % of cell counts |
| 10D0S | Blastula | 11-20um | 14.879 |
| 10D0S | Blastula | 21-30um | 35.772 |
| 10D0S | Blastula | 31-40um | 32.505 |
| 10D0S | Blastula | 41-50um | 11.878 |
| 10D0S | Blastula | 51-60um | 5.535 |
| 10D0S | Blastula | 61-70um | 5.626 |
| 10D0S | Blastula | 71-80um | 3.505 |
| 10D1S | Blastula | 0-10um | 8.437 |
| 10D1S | Blastula | 11-20um | 18.428 |
| 10D1S | Blastula | 21-30um | 33.213 |
| 10D1S | Blastula | 31-40um | 23.036 |
| 10D1S | Blastula | 41-50um | 17.401 |
| 10D1S | Blastula | 51-60um | 8.892 |
| 10D1S | Blastula | 61-70um | 7.569 |
| 10D1S | Blastula | 71-80um | 5.669 |
| 10D10S | Blastula | 11-20um | 8.512 |
| 10D10S | Blastula | 21-30um | 17.044 |
| 10D10S | Blastula | 31-40um | 25.338 |
| 10D10S | Blastula | 41-50um | 22.254 |
| 10D10S | Blastula | 51-60um | 14.293 |
| 10D10S | Blastula | 61-70um | 11.16 |
| 10D10S | Blastula | 71-80um | 11.673 |
| 10D10S | Blastula | 81-90um | 7.316 |
| 10D10S | Blastula | 91-100um | 5.225 |
| 10D10S | Blastula | 101-110um | 6.031 |
| Pre-freeze | Blastula | 0-10um | 1.373 |
| Pre-freeze | Blastula | 11-20um | 0.857 |
| Pre-freeze | Blastula | 21-30um | 4.466 |
| Pre-freeze | Blastula | 31-40um | 21.206 |
| Pre-freeze | Blastula | 41-50um | 32.715 |
| Pre-freeze | Blastula | 51-60um | 20.916 |
| Pre-freeze | Blastula | 61-70um | 7.929 |
| Pre-freeze | Blastula | 71-80um | 13.811 |
| Pre-freeze | Blastula | 81-90um | 2.128 |
| Pre-freeze | Blastula | 91-100um | 1.19 |
| Pre-freeze | Blastula | 101-110um | 1.525 |
| 10D0S | gastrula | 0-10um | 4.941 |
| 10D0S | gastrula | 11-20um | 29.995 |
| 10D0S | gastrula | 21-30um | 47.222 |
| 10D0S | gastrula | 31-40um | 16.498 |
| 10D0S | gastrula | 41-50um | 7.49 |
| 10D0S | gastrula | 51-60um | 4.7 |
| 10D0S | gastrula | 71-80um | 3.19 |
| 10D1S | gastrula | 0-10um | 2.598 |
| 10D1S | gastrula | 11-20um | 21.953 |
| 10D1S | gastrula | 21-30um | 51.024 |
| 10D1S | gastrula | 31-40um | 16.624 |
| 10D1S | gastrula | 41-50um | 8.561 |
| 10D1S | gastrula | 51-60um | 2.707 |
| 10D1S | gastrula | 61-70um | 3.378 |
| 10D1S | gastrula | 71-80um | 5.052 |
| 10D10S | gastrula | 0-10um | 1.324 |
| 10D10S | gastrula | 11-20um | 14.01 |
| 10D10S | gastrula | 21-30um | 65.415 |
| 10D10S | gastrula | 31-40um | 16.255 |
| 10D10S | gastrula | 41-50um | 3.796 |
| 10D10S | gastrula | 51-60um | 2.285 |
| 10D10S | gastrula | 61-70um | 3.575 |
| 10D10S | gastrula | 71-80um | 1.33 |
| Pre-freeze | gastrula | 0-10um | 0.413 |
| Pre-freeze | gastrula | 11-20um | 22.616 |
| Pre-freeze | gastrula | 21-30um | 55.11 |
| Pre-freeze | gastrula | 31-40um | 10.672 |
| Pre-freeze | gastrula | 41-50um | 3.834 |
| Pre-freeze | gastrula | 51-60um | 0.504 |
| 10D0S | neurula | 0-10um | 3.003 |
| 10D0S | neurula | 11-20um | 34.414 |
| 10D0S | neurula | 21-30um | 46.07 |
| 10D0S | neurula | 31-40um | 12.988 |
| 10D0S | neurula | 41-50um | 5.223 |
| 10D0S | neurula | 51-60um | 2.326 |
| 10D1S | neurula | 0-10um | 3.114 |
| 10D1S | neurula | 11-20um | 28.941 |
| 10D1S | neurula | 21-30um | 55.69 |
| 10D1S | neurula | 31-40um | 10.285 |
| 10D1S | neurula | 41-50um | 3.411 |
| 10D1S | neurula | 51-60um | 2.636 |
| 10D1S | neurula | 111-120um | 2.569 |
| 10D10S | neurula | 0-10um | 1.168 |
| 10D10S | neurula | 11-20um | 30.188 |
| 10D10S | neurula | 21-30um | 58.38 |
| 10D10S | neurula | 31-40um | 7.013 |
| 10D10S | neurula | 41-50um | 2.866 |
| 10D10S | neurula | 51-60um | 1.234 |
| 10D10S | neurula | 71-80um | 0.567 |
| Pre-freeze | neurula | 0-10um | 0.508 |
| Pre-freeze | neurula | 11-20um | 62.094 |
| Pre-freeze | neurula | 21-30um | 30.371 |
| Pre-freeze | neurula | 31-40um | 5.189 |
| Pre-freeze | neurula | 41-50um | 1.41 |

Supplementary Table 5: Odds ratios and 95% confidence intervals for pairwise comparisons of size class distributions between cryoprotectant levels for each stage (blastula, gastrula, and neurula) in *L. peronii.*

| Size Class (µm) Distributions | | | | | | | |
| --- | --- | --- | --- | --- | --- | --- | --- |
| reverse contrast | Odds ratio | LCL^1^ | UCL^2^ | contrast | odds ratio | LCL^1^ | UCL^2^ |
| (pre-freeze 0-10um blast) / (10D1S 0-10um blast) | 0.151 | 0.008 | 2.881 | (10D1S 0-10um blast) / (pre-freeze 0-10um blast) | 6.621 | 0.347 | 126.293 |
| (10D1S 0-10um gast) / (10D0S 0-10um gast) | 0.513 | 0.069 | 3.824 | (10D0S 0-10um gast) / (10D1S 0-10um gast) | 1.949 | 0.261 | 14.520 |
| (10D10S 0-10um gast) / (10D0S 0-10um gast) | 0.258 | 0.034 | 1.980 | (10D0S 0-10um gast) / (10D10S 0-10um gast) | 3.874 | 0.505 | 29.715 |
| (10D10S 0-10um gast) / (10D1S 0-10um gast) | 0.503 | 0.056 | 4.533 | (10D0S 0-10um gast) / (pre-freeze 0-10um gast) | 12.526* | 1.351 | 116.134 |
| (pre-freeze 0-10um gast) / (10D0S 0-10um gast) | 0.080* | 0.009 | 0.740 | (10D1S 0-10um gast) / (10D10S 0-10um gast) | 1.988 | 0.221 | 17.915 |
| (pre-freeze 0-10um gast) / (10D1S 0-10um gast) | 0.156 | 0.014 | 1.673 | (10D1S 0-10um gast) / (pre-freeze 0-10um gast) | 6.428 | 0.598 | 69.127 |
| (pre-freeze 0-10um gast) / (10D10S 0-10um gast) | 0.309 | 0.028 | 3.408 | (10D10S 0-10um gast) / (pre-freeze 0-10um gast) | 3.233 | 0.293 | 35.633 |
| (10D1S 0-10um neur) / (10D0S 0-10um neur) | 1.038 | 0.291 | 3.699 | (10D0S 0-10um neur) / (10D1S 0-10um neur) | 0.963 | 0.270 | 3.432 |
| (10D10S 0-10um neur) / (10D0S 0-10um neur) | 0.382 | 0.100 | 1.460 | (10D0S 0-10um neur) / (10D10S 0-10um neur) | 2.620 | 0.685 | 10.023 |
| (10D10S 0-10um neur) / (10D1S 0-10um neur) | 0.368* | 0.137 | 0.985 | (10D0S 0-10um neur) / (pre-freeze 0-10um neur) | 6.058* | 1.419 | 25.857 |
| (pre-freeze 0-10um neur) / (10D0S 0-10um neur) | 0.165* | 0.039 | 0.705 | (10D1S 0-10um neur) / (10D10S 0-10um neur) | 2.720* | 1.016 | 7.286 |
| (pre-freeze 0-10um neur) / (10D1S 0-10um neur) | 0.159* | 0.051 | 0.492 | (10D1S 0-10um neur) / (pre-freeze 0-10um neur) | 6.289* | 2.032 | 19.468 |
| (pre-freeze 0-10um neur) / (10D10S 0-10um neur) | 0.433 | 0.129 | 1.449 | (10D10S 0-10um neur) / (pre-freeze 0-10um neur) | 2.312 | 0.690 | 7.748 |
| (10D1S 11-20um blast) / (10D0S 11-20um blast) | 1.292 | 0.724 | 2.307 | (10D0S 11-20um blast) / (10D1S 11-20um blast) | 0.774 | 0.433 | 1.381 |
| (10D10S 11-20um blast) / (10D0S 11-20um blast) | 0.532* | 0.287 | 0.986 | (10D0S 11-20um blast) / (10D10S 11-20um blast) | 1.879* | 1.015 | 3.479 |
| (10D10S 11-20um blast) / (10D1S 11-20um blast) | 0.412* | 0.225 | 0.753 | (10D0S 11-20um blast) / (pre-freeze 11-20um blast) | 20.214* | 1.536 | 266.004 |
| (pre-freeze 11-20um blast) / (10D0S 11-20um blast) | 0.049* | 0.004 | 0.651 | (10D1S 11-20um blast) / (10D10S 11-20um blast) | 2.428* | 1.328 | 4.438 |
| (pre-freeze 11-20um blast) / (10D1S 11-20um blast) | 0.038* | 0.003 | 0.502 | (10D1S 11-20um blast) / (pre-freeze 11-20um blast) | 26.124* | 1.991 | 342.743 |
| (pre-freeze 11-20um blast) / (10D10S 11-20um blast) | 0.093 | 0.007 | 1.230 | (10D10S 11-20um blast) / (pre-freeze 11-20um blast) | 10.759 | 0.813 | 142.363 |
| (10D1S 11-20um gast) / (10D0S 11-20um gast) | 0.656 | 0.429 | 1.005 | (10D0S 11-20um gast) / (10D1S 11-20um gast) | 1.523 | 0.995 | 2.331 |
| (10D10S 11-20um gast) / (10D0S 11-20um gast) | 0.380* | 0.250 | 0.579 | (10D0S 11-20um gast) / (10D10S 11-20um gast) | 2.630* | 1.727 | 4.003 |
| (10D10S 11-20um gast) / (10D1S 11-20um gast) | 0.579* | 0.380 | 0.883 | (10D0S 11-20um gast) / (pre-freeze 11-20um gast) | 1.466 | 0.804 | 2.672 |
| (pre-freeze 11-20um gast) / (10D0S 11-20um gast) | 0.682 | 0.374 | 1.243 | (10D1S 11-20um gast) / (10D10S 11-20um gast) | 1.726* | 1.133 | 2.631 |
| (pre-freeze 11-20um gast) / (10D1S 11-20um gast) | 1.039 | 0.570 | 1.896 | (10D1S 11-20um gast) / (pre-freeze 11-20um gast) | 0.962 | 0.528 | 1.756 |
| (pre-freeze 11-20um gast) / (10D10S 11-20um gast) | 1.794 | 0.986 | 3.263 | (10D10S 11-20um gast) / (pre-freeze 11-20um gast) | 0.557 | 0.307 | 1.014 |
| (10D1S 11-20um neur) / (10D0S 11-20um neur) | 0.776 | 0.531 | 1.134 | (10D0S 11-20um neur) / (10D1S 11-20um neur) | 1.288 | 0.882 | 1.882 |
| (10D10S 11-20um neur) / (10D0S 11-20um neur) | 0.824 | 0.571 | 1.189 | (10D0S 11-20um neur) / (10D10S 11-20um neur) | 1.213 | 0.841 | 1.751 |
| (10D10S 11-20um neur) / (10D1S 11-20um neur) | 1.062 | 0.740 | 1.523 | (10D0S 11-20um neur) / (pre-freeze 11-20um neur) | 0.320* | 0.185 | 0.555 |
| (pre-freeze 11-20um neur) / (10D0S 11-20um neur) | 3.122* | 1.801 | 5.411 | (10D1S 11-20um neur) / (10D10S 11-20um neur) | 0.942 | 0.656 | 1.351 |
| (pre-freeze 11-20um neur) / (10D1S 11-20um neur) | 4.022* | 2.330 | 6.944 | (10D1S 11-20um neur) / (pre-freeze 11-20um neur) | 0.249* | 0.144 | 0.429 |
| (pre-freeze 11-20um neur) / (10D10S 11-20um neur) | 3.788* | 2.213 | 6.486 | (10D10S 11-20um neur) / (pre-freeze 11-20um neur) | 0.264* | 0.154 | 0.452 |
| (10D1S 21-30um blast) / (10D0S 21-30um blast) | 0.893 | 0.556 | 1.433 | (10D0S 21-30um blast) / (10D1S 21-30um blast) | 1.120 | 0.698 | 1.798 |
| (10D10S 21-30um blast) / (10D0S 21-30um blast) | 0.369* | 0.225 | 0.604 | (10D0S 21-30um blast) / (10D10S 21-30um blast) | 2.711* | 1.654 | 4.442 |
| (10D10S 21-30um blast) / (10D1S 21-30um blast) | 0.413* | 0.252 | 0.676 | (10D0S 21-30um blast) / (pre-freeze 21-30um blast) | 11.915* | 5.009 | 28.347 |
| (pre-freeze 21-30um blast) / (10D0S 21-30um blast) | 0.084* | 0.035 | 0.200 | (10D1S 21-30um blast) / (10D10S 21-30um blast) | 2.420* | 1.479 | 3.961 |
| (pre-freeze 21-30um blast) / (10D1S 21-30um blast) | 0.094* | 0.040 | 0.223 | (10D1S 21-30um blast) / (pre-freeze 21-30um blast) | 10.639* | 4.475 | 25.292 |
| (pre-freeze 21-30um blast) / (10D10S 21-30um blast) | 0.227* | 0.095 | 0.547 | (10D10S 21-30um blast) / (pre-freeze 21-30um blast) | 4.396* | 1.828 | 10.570 |
| (10D1S 21-30um gast) / (10D0S 21-30um gast) | 1.164 | 0.781 | 1.736 | (10D0S 21-30um gast) / (10D1S 21-30um gast) | 0.859 | 0.576 | 1.280 |
| (10D10S 21-30um gast) / (10D0S 21-30um gast) | 2.114* | 1.441 | 3.102 | (10D0S 21-30um gast) / (10D10S 21-30um gast) | 0.473* | 0.322 | 0.694 |
| (10D10S 21-30um gast) / (10D1S 21-30um gast) | 1.815* | 1.247 | 2.643 | (10D0S 21-30um gast) / (pre-freeze 21-30um gast) | 0.729 | 0.412 | 1.290 |
| (pre-freeze 21-30um gast) / (10D0S 21-30um gast) | 1.372 | 0.775 | 2.429 | (10D1S 21-30um gast) / (10D10S 21-30um gast) | 0.551* | 0.378 | 0.802 |
| (pre-freeze 21-30um gast) / (10D1S 21-30um gast) | 1.178 | 0.669 | 2.075 | (10D1S 21-30um gast) / (pre-freeze 21-30um gast) | 0.849 | 0.482 | 1.494 |
| (pre-freeze 21-30um gast) / (10D10S 21-30um gast) | 0.649 | 0.373 | 1.131 | (10D10S 21-30um gast) / (pre-freeze 21-30um gast) | 1.541 | 0.885 | 2.683 |
| (10D1S 21-30um neur) / (10D0S 21-30um neur) | 1.471* | 1.017 | 2.128 | (10D0S 21-30um neur) / (10D1S 21-30um neur) | 0.680* | 0.470 | 0.983 |
| (10D10S 21-30um neur) / (10D0S 21-30um neur) | 1.642* | 1.146 | 2.352 | (10D0S 21-30um neur) / (10D10S 21-30um neur) | 0.609* | 0.425 | 0.872 |
| (10D10S 21-30um neur) / (10D1S 21-30um neur) | 1.116 | 0.785 | 1.586 | (10D0S 21-30um neur) / (pre-freeze 21-30um neur) | 1.959* | 1.131 | 3.390 |
| (pre-freeze 21-30um neur) / (10D0S 21-30um neur) | 0.511* | 0.295 | 0.884 | (10D1S 21-30um neur) / (10D10S 21-30um neur) | 0.896 | 0.630 | 1.273 |
| (pre-freeze 21-30um neur) / (10D1S 21-30um neur) | 0.347* | 0.202 | 0.598 | (10D1S 21-30um neur) / (pre-freeze 21-30um neur) | 2.881 | 1.673 | 4.963 |
| (pre-freeze 21-30um neur) / (10D10S 21-30um neur) | 0.311* | 0.182 | 0.532 | (10D10S 21-30um neur) / (pre-freeze 21-30um neur) | 3.216 | 1.880 | 5.502 |
| (10D1S 31-40um blast) / (10D0S 31-40um blast) | 0.621 | 0.377 | 1.025 | (10D0S 31-40um blast) / (10D1S 31-40um blast) | 1.609 | 0.976 | 2.654 |
| (10D10S 31-40um blast) / (10D0S 31-40um blast) | 0.705 | 0.434 | 1.144 | (10D0S 31-40um blast) / (10D10S 31-40um blast) | 1.419 | 0.874 | 2.303 |
| (10D10S 31-40um blast) / (10D1S 31-40um blast) | 1.134 | 0.685 | 1.876 | (10D0S 31-40um blast) / (pre-freeze 31-40um blast) | 1.789 | 0.922 | 3.474 |
| (pre-freeze 31-40um blast) / (10D0S 31-40um blast) | 0.559 | 0.288 | 1.085 | (10D1S 31-40um blast) / (10D10S 31-40um blast) | 0.882 | 0.533 | 1.460 |
| (pre-freeze 31-40um blast) / (10D1S 31-40um blast) | 0.899 | 0.457 | 1.771 | (10D1S 31-40um blast) / (pre-freeze 31-40um blast) | 1.112 | 0.565 | 2.190 |
| (pre-freeze 31-40um blast) / (10D10S 31-40um blast) | 0.793 | 0.408 | 1.543 | (10D10S 31-40um blast) / (pre-freeze 31-40um blast) | 1.261 | 0.648 | 2.454 |
| (10D1S 31-40um gast) / (10D0S 31-40um gast) | 1.009 | 0.636 | 1.601 | (10D0S 31-40um gast) / (10D1S 31-40um gast) | 0.991 | 0.625 | 1.572 |
| (10D10S 31-40um gast) / (10D0S 31-40um gast) | 0.982 | 0.635 | 1.519 | (10D0S 31-40um gast) / (10D10S 31-40um gast) | 1.018 | 0.658 | 1.574 |
| (10D10S 31-40um gast) / (10D1S 31-40um gast) | 0.973 | 0.637 | 1.487 | (10D0S 31-40um gast) / (pre-freeze 31-40um gast) | 1.654 | 0.877 | 3.119 |
| (pre-freeze 31-40um gast) / (10D0S 31-40um gast) | 0.605 | 0.321 | 1.140 | (10D1S 31-40um gast) / (10D10S 31-40um gast) | 1.027 | 0.672 | 1.569 |
| (pre-freeze 31-40um gast) / (10D1S 31-40um gast) | 0.599 | 0.320 | 1.121 | (10D1S 31-40um gast) / (pre-freeze 31-40um gast) | 1.669 | 0.892 | 3.122 |
| (pre-freeze 31-40um gast) / (10D10S 31-40um gast) | 0.616 | 0.335 | 1.129 | (10D10S 31-40um gast) / (pre-freeze 31-40um gast) | 1.625 | 0.885 | 2.981 |
| (10D1S 31-40um neur) / (10D0S 31-40um neur) | 0.768 | 0.486 | 1.214 | (10D0S 31-40um neur) / (10D1S 31-40um neur) | 1.302 | 0.824 | 2.058 |
| (10D10S 31-40um neur) / (10D0S 31-40um neur) | 0.505* | 0.323 | 0.789 | (10D0S 31-40um neur) / (10D10S 31-40um neur) | 1.979* | 1.267 | 3.093 |
| (10D10S 31-40um neur) / (10D1S 31-40um neur) | 0.658 | 0.419 | 1.032 | (10D0S 31-40um neur) / (pre-freeze 31-40um neur) | 2.728* | 1.460 | 5.095 |
| (pre-freeze 31-40um neur) / (10D0S 31-40um neur) | 0.367* | 0.196 | 0.685 | (10D1S 31-40um neur) / (10D10S 31-40um neur) | 1.520 | 0.969 | 2.384 |
| (pre-freeze 31-40um neur) / (10D1S 31-40um neur) | 0.477* | 0.255 | 0.894 | (10D1S 31-40um neur) / (pre-freeze 31-40um neur) | 2.095* | 1.118 | 3.925 |
| (pre-freeze 31-40um neur) / (10D10S 31-40um neur) | 0.726 | 0.390 | 1.348 | (10D10S 31-40um neur) / (pre-freeze 31-40um neur) | 1.378 | 0.742 | 2.561 |
| (10D1S 41-50um blast) / (10D0S 41-50um blast) | 1.563 | 0.850 | 2.874 | (10D0S 41-50um blast) / (10D1S 41-50um blast) | 0.640 | 0.348 | 1.177 |
| (10D10S 41-50um blast) / (10D0S 41-50um blast) | 2.124* | 1.188 | 3.796 | (10D0S 41-50um blast) / (10D10S 41-50um blast) | 0.471* | 0.263 | 0.842 |
| (10D10S 41-50um blast) / (10D1S 41-50um blast) | 1.359 | 0.779 | 2.371 | (10D0S 41-50um blast) / (pre-freeze 41-50um blast) | 0.277* | 0.137 | 0.563 |
| (pre-freeze 41-50um blast) / (10D0S 41-50um blast) | 3.607* | 1.778 | 7.319 | (10D1S 41-50um blast) / (10D10S 41-50um blast) | 0.736 | 0.422 | 1.285 |
| (pre-freeze 41-50um blast) / (10D1S 41-50um blast) | 2.308* | 1.160 | 4.590 | (10D1S 41-50um blast) / (pre-freeze 41-50um blast) | 0.433* | 0.218 | 0.862 |
| (pre-freeze 41-50um blast) / (10D10S 41-50um blast) | 1.699 | 0.875 | 3.296 | (10D10S 41-50um blast) / (pre-freeze 41-50um blast) | 0.589 | 0.303 | 1.142 |
| (10D1S 41-50um gast) / (10D0S 41-50um gast) | 1.156 | 0.533 | 2.510 | (10D0S 41-50um gast) / (10D1S 41-50um gast) | 0.865 | 0.398 | 1.878 |
| (10D10S 41-50um gast) / (10D0S 41-50um gast) | 0.487 | 0.215 | 1.105 | (10D0S 41-50um gast) / (10D10S 41-50um gast) | 2.052 | 0.905 | 4.654 |
| (10D10S 41-50um gast) / (10D1S 41-50um gast) | 0.422* | 0.210 | 0.848 | (10D0S 41-50um gast) / (pre-freeze 41-50um gast) | 2.031 | 0.834 | 4.946 |
| (pre-freeze 41-50um gast) / (10D0S 41-50um gast) | 0.492 | 0.202 | 1.199 | (10D1S 41-50um gast) / (10D10S 41-50um gast) | 2.372* | 1.180 | 4.771 |
| (pre-freeze 41-50um gast) / (10D1S 41-50um gast) | 0.426* | 0.195 | 0.930 | (10D1S 41-50um gast) / (pre-freeze 41-50um gast) | 2.348* | 1.075 | 5.129 |
| (pre-freeze 41-50um gast) / (10D10S 41-50um gast) | 1.010 | 0.443 | 2.304 | (10D10S 41-50um gast) / (pre-freeze 41-50um gast) | 0.990 | 0.434 | 2.258 |
| (10D1S 41-50um neur) / (10D0S 41-50um neur) | 0.641 | 0.307 | 1.336 | (10D0S 41-50um neur) / (10D1S 41-50um neur) | 1.560 | 0.748 | 3.253 |
| (10D10S 41-50um neur) / (10D0S 41-50um neur) | 0.535 | 0.267 | 1.074 | (10D0S 41-50um neur) / (10D10S 41-50um neur) | 1.868 | 0.931 | 3.748 |
| (10D10S 41-50um neur) / (10D1S 41-50um neur) | 0.835 | 0.395 | 1.764 | (10D0S 41-50um neur) / (pre-freeze 41-50um neur) | 3.854* | 1.670 | 8.893 |
| (pre-freeze 41-50um neur) / (10D0S 41-50um neur) | 0.260* | 0.112 | 0.599 | (10D1S 41-50um neur) / (10D10S 41-50um neur) | 1.197 | 0.567 | 2.529 |
| (pre-freeze 41-50um neur) / (10D1S 41-50um neur) | 0.405* | 0.168 | 0.976 | (10D1S 41-50um neur) / (pre-freeze 41-50um neur) | 2.470* | 1.025 | 5.951 |
| (pre-freeze 41-50um neur) / (10D10S 41-50um neur) | 0.485 | 0.208 | 1.132 | (10D10S 41-50um neur) / (pre-freeze 41-50um neur) | 2.063 | 0.884 | 4.816 |
| (10D1S 51-60um blast) / (10D0S 51-60um blast) | 1.666 | 0.526 | 5.275 | (10D0S 51-60um blast) / (10D1S 51-60um blast) | 0.600 | 0.190 | 1.902 |
| (10D10S 51-60um blast) / (10D0S 51-60um blast) | 2.846 | 0.967 | 8.379 | (10D0S 51-60um blast) / (10D10S 51-60um blast) | 0.351 | 0.119 | 1.034 |
| (10D10S 51-60um blast) / (10D1S 51-60um blast) | 1.709 | 0.869 | 3.360 | (10D0S 51-60um blast) / (pre-freeze 51-60um blast) | 0.222* | 0.069 | 0.706 |
| (pre-freeze 51-60um blast) / (10D0S 51-60um blast) | 4.514* | 1.416 | 14.389 | (10D1S 51-60um blast) / (10D10S 51-60um blast) | 0.585 | 0.298 | 1.151 |
| (pre-freeze 51-60um blast) / (10D1S 51-60um blast) | 2.710* | 1.221 | 6.012 | (10D1S 51-60um blast) / (pre-freeze 51-60um blast) | 0.369* | 0.166 | 0.819 |
| (pre-freeze 51-60um blast) / (10D10S 51-60um blast) | 1.586 | 0.798 | 3.152 | (10D10S 51-60um blast) / (pre-freeze 51-60um blast) | 0.631 | 0.317 | 1.253 |
| (10D1S 51-60um gast) / (10D0S 51-60um gast) | 0.564 | 0.071 | 4.475 | (10D0S 51-60um gast) / (10D1S 51-60um gast) | 1.773 | 0.223 | 14.063 |
| (10D10S 51-60um gast) / (10D0S 51-60um gast) | 0.474 | 0.068 | 3.293 | (10D0S 51-60um gast) / (10D10S 51-60um gast) | 2.109 | 0.304 | 14.650 |
| (10D10S 51-60um gast) / (10D1S 51-60um gast) | 0.840 | 0.239 | 2.960 | (10D0S 51-60um gast) / (pre-freeze 51-60um gast) | 9.731* | 1.061 | 89.204 |
| (pre-freeze 51-60um gast) / (10D0S 51-60um gast) | 0.103* | 0.011 | 0.942 | (10D1S 51-60um gast) / (10D10S 51-60um gast) | 1.190 | 0.338 | 4.191 |
| (pre-freeze 51-60um gast) / (10D1S 51-60um gast) | 0.182* | 0.035 | 0.953 | (10D1S 51-60um gast) / (pre-freeze 51-60um gast) | 5.489* | 1.049 | 28.723 |
| (pre-freeze 51-60um gast) / (10D10S 51-60um gast) | 0.217* | 0.049 | 0.957 | (10D10S 51-60um gast) / (pre-freeze 51-60um gast) | 4.613* | 1.045 | 20.373 |
| (10D1S 51-60um neur) / (10D0S 51-60um neur) | 1.137 | 0.136 | 9.507 | (10D0S 51-60um neur) / (10D1S 51-60um neur) | 0.880 | 0.105 | 7.354 |
| (10D10S 51-60um neur) / (10D0S 51-60um neur) | 0.524 | 0.118 | 2.329 | (10D0S 51-60um neur) / (10D10S 51-60um neur) | 1.907 | 0.429 | 8.469 |
| (10D10S 51-60um neur) / (10D1S 51-60um neur) | 0.461 | 0.069 | 3.065 | (10D0S 51-60um neur) / (pre-freeze 51-60um neur) | NA | NA | NA |
| (pre-freeze 51-60um neur) / (10D0S 51-60um neur) | NA | NA | NA | (10D1S 51-60um neur) / (10D10S 51-60um neur) | 2.168 | 0.326 | 14.404 |
| (10D1S 61-70um blast) / (10D0S 61-70um blast) | 1.374 | 0.397 | 4.752 | (10D0S 61-70um blast) / (10D1S 61-70um blast) | 0.728 | 0.210 | 2.519 |
| (10D10S 61-70um blast) / (10D0S 61-70um blast) | 2.107 | 0.715 | 6.213 | (10D0S 61-70um blast) / (10D10S 61-70um blast) | 0.475 | 0.161 | 1.399 |
| (10D10S 61-70um blast) / (10D1S 61-70um blast) | 1.534 | 0.602 | 3.910 | (10D0S 61-70um blast) / (pre-freeze 61-70um blast) | 0.692 | 0.214 | 2.243 |
| (pre-freeze 61-70um blast) / (10D0S 61-70um blast) | 1.445 | 0.446 | 4.680 | (10D1S 61-70um blast) / (10D10S 61-70um blast) | 0.652 | 0.256 | 1.662 |
| (pre-freeze 61-70um blast) / (10D1S 61-70um blast) | 1.052 | 0.371 | 2.984 | (10D1S 61-70um blast) / (pre-freeze 61-70um blast) | 0.951 | 0.335 | 2.699 |
| (pre-freeze 61-70um blast) / (10D10S 61-70um blast) | 0.686 | 0.294 | 1.597 | (10D10S 61-70um blast) / (pre-freeze 61-70um blast) | 1.459 | 0.626 | 3.399 |
| (10D10S 61-70um gast) / (10D1S 61-70um gast) | 1.060 | 0.057 | 19.734 | (10D0S 61-70um gast) / (pre-freeze 61-70um gast) | NA | NA | NA |
| (pre-freeze 61-70um gast) / (10D0S 61-70um gast) | NA | NA | NA | (10D1S 61-70um gast) / (10D10S 61-70um gast) | 0.943 | 0.051 | 17.552 |
| (10D1S 71-80um blast) / (10D0S 71-80um blast) | 1.655 | 0.098 | 27.911 | (10D0S 71-80um blast) / (10D1S 71-80um blast) | 0.604 | 0.036 | 10.196 |
| (10D10S 71-80um blast) / (10D0S 71-80um blast) | 3.638 | 0.317 | 41.820 | (10D0S 71-80um blast) / (10D10S 71-80um blast) | 0.275 | 0.024 | 3.159 |
| (10D10S 71-80um blast) / (10D1S 71-80um blast) | 2.199 | 0.427 | 11.319 | (10D0S 71-80um blast) / (pre-freeze 71-80um blast) | 0.227 | 0.017 | 3.001 |
| (pre-freeze 71-80um blast) / (10D0S 71-80um blast) | 4.412 | 0.333 | 58.419 | (10D1S 71-80um blast) / (10D10S 71-80um blast) | 0.455 | 0.088 | 2.341 |
| (pre-freeze 71-80um blast) / (10D1S 71-80um blast) | 2.667 | 0.422 | 16.835 | (10D1S 71-80um blast) / (pre-freeze 71-80um blast) | 0.375 | 0.059 | 2.368 |
| (pre-freeze 71-80um blast) / (10D10S 71-80um blast) | 1.213 | 0.375 | 3.916 | (10D10S 71-80um blast) / (pre-freeze 71-80um blast) | 0.825 | 0.255 | 2.663 |
| (10D1S 71-80um gast) / (10D0S 71-80um gast) | 1.615 | 0.072 | 36.465 | (10D0S 71-80um gast) / (10D1S 71-80um gast) | 0.619 | 0.027 | 13.983 |
| (10D10S 71-80um gast) / (10D0S 71-80um gast) | 0.409 | 0.029 | 5.797 | (10D0S 71-80um gast) / (10D10S 71-80um gast) | 2.445 | 0.172 | 34.661 |
| (10D10S 71-80um gast) / (10D1S 71-80um gast) | 0.253 | 0.018 | 3.619 | (10D0S 71-80um gast) / (pre-freeze 71-80um gast) | NA | NA | NA |
| (pre-freeze 81-90um blast) / (10D10S 81-90um blast) | 0.275 | 0.033 | 2.288 | (10D1S 71-80um gast) / (10D10S 71-80um gast) | 3.949 | 0.276 | 56.423 |
| (pre-freeze 91-100um blast) / (10D10S 91-100um blast) | 0.219 | 0.018 | 2.610 | (10D10S 81-90um blast) / (pre-freeze 81-90um blast) | 3.630 | 0.437 | 30.161 |
| (pre-freeze 101-110um blast) / (10D10S 101-110um blast) | 0.241 | 0.018 | 3.318 | (10D10S 91-100um blast) / (pre-freeze 91-100um blast) | 4.576 | 0.383 | 54.642 |

^1^Lower confidence limit, ^2^Upper confidence limit, * Significant pairwise comparisons

Supplementary Table 8: *R. littlejohni* (EMMs) for percentage of neurula cells in each size class for pre-freeze vs. 10D10S.

| treatment | size class (µm) | % cell counts |
| --- | --- | --- |
| 10D10S | 0-10um | 1.494 |
| 10D10S | 11-20um | 19.358 |
| 10D10S | 21-30um | 58.773 |
| 10D10S | 31-40um | 14.507 |
| 10D10S | 41-50um | 4.234 |
| 10D10S | 51-60um | 2.746 |
| 10D10S | 61-70um | 1.634 |
| 10D10S | 71-80um | 1.208 |
| 10D10S | 81-90um | 1.714 |
| 10D10S | 101-110um | 1.849 |
| Pre-freeze | 0-10um | 0.347 |
| Pre-freeze | 11-20um | 34.104 |
| Pre-freeze | 21-30um | 52.917 |
| Pre-freeze | 31-40um | 7.495 |
| Pre-freeze | 41-50um | 1.431 |
| Pre-freeze | 51-60um | 1.033 |
| Pre-freeze | 61-70um | 0.379 |
| Pre-freeze | 71-80um | 0.293 |
| Pre-freeze | 101-110um | 0.420 |
| Pre-freeze | 141-150um | 0.293 |

Supplementary Table 9: Odds ratios and 95% confidence intervals for pairwise comparisons of percentages within size class and cryoprotectant combinations for *R. littlejohni* neurula cells (pre-freeze vs. 10D10S).

| LLJ Size class | | | | | | | |
| --- | --- | --- | --- | --- | --- | --- | --- |
| reverse contrast | odds ratio | LCL^1^ | UCL^2^ | contrast | odds ratio | LCL^1^ | UCL^2^ |
| (pre 0-10um) / (10D10S 0-10um) | 0.229 | 0.033 | 1.579 | (10D10S 0-10um) / (pre 0-10um) | 4.361 | 0.633 | 30.036 |
| (10D10S 11-20um) / (10D10S 0-10um) | 15.83* | 4.454 | 56.265 | (10D10S 0-10um) / (10D10S 11-20um) | 0.063* | 0.018 | 0.225 |
| (10D10S 11-20um) / (pre 0-10um) | 69.033* | 14.99 | 317.919 | (10D10S 0-10um) / (pre 11-20um) | 0.029* | 0.008 | 0.108 |
| (pre 11-20um) / (10D10S 0-10um) | 34.129* | 9.232 | 126.169 | (10D10S 0-10um) / (10D10S 21-30um) | 0.011* | 0.003 | 0.038 |
| (pre 11-20um) / (pre 0-10um) | 148.83* | 31.274 | 708.279 | (10D10S 0-10um) / (pre 21-30um) | 0.013* | 0.004 | 0.05 |
| (pre 11-20um) / (10D10S 11-20um) | 2.156* | 1.227 | 3.789 | (10D10S 0-10um) / (10D10S 31-40um) | 0.089* | 0.025 | 0.319 |
| (10D10S 21-30um) / (10D10S 0-10um) | 94.012* | 26.605 | 332.198 | (10D10S 0-10um) / (pre 31-40um) | 0.187* | 0.05 | 0.706 |
| (10D10S 21-30um) / (pre 0-10um) | 409.967* | 89.451 | 1878.932 | (10D10S 0-10um) / (10D10S 41-50um) | 0.343 | 0.09 | 1.307 |
| (10D10S 21-30um) / (10D10S 11-20um) | 5.939* | 3.79 | 9.305 | (10D10S 0-10um) / (pre 41-50um) | 1.044 | 0.25 | 4.365 |
| (10D10S 21-30um) / (pre 11-20um) | 2.755* | 1.591 | 4.77 | (10D10S 0-10um) / (10D10S 51-60um) | 0.537 | 0.116 | 2.495 |
| (pre 21-30um) / (10D10S 0-10um) | 74.116* | 20.071 | 273.693 | (10D10S 0-10um) / (pre 51-60um) | 1.453 | 0.332 | 6.355 |
| (pre 21-30um) / (pre 0-10um) | 323.206* | 67.977 | 1536.715 | (10D10S 0-10um) / (10D10S 61-70um) | 0.913 | 0.131 | 6.362 |
| (pre 21-30um) / (10D10S 11-20um) | 4.682* | 2.671 | 8.206 | (10D10S 0-10um) / (pre 61-70um) | 3.984 | 0.708 | 22.414 |
| (pre 21-30um) / (pre 11-20um) | 2.172* | 1.14 | 4.135 | (10D10S 0-10um) / (10D10S 71-80um) | 1.24 | 0.108 | 14.293 |
| (pre 21-30um) / (10D10S 21-30um) | 0.788 | 0.457 | 1.361 | (10D10S 0-10um) / (pre 71-80um) | 5.16 | 0.452 | 58.961 |
| (10D10S 31-40um) / (10D10S 0-10um) | 11.19* | 3.134 | 39.952 | (10D10S 0-10um) / (10D10S 81-90um) | 0.869 | 0.075 | 10.065 |
| (10D10S 31-40um) / (pre 0-10um) | 48.796* | 10.555 | 225.572 | (10D10S 0-10um) / (10D10S 101-110um) | 0.805 | 0.069 | 9.33 |
| (10D10S 31-40um) / (10D10S 11-20um) | 0.707 | 0.438 | 1.14 | (10D10S 0-10um) / (pre 101-110um) | 3.595 | 0.314 | 41.129 |
| (10D10S 31-40um) / (pre 11-20um) | 0.328* | 0.185 | 0.582 | (10D10S 0-10um) / (pre 141-150um) | 5.16 | 0.452 | 58.961 |
| (10D10S 31-40um) / (10D10S 21-30um) | 0.119* | 0.075 | 0.189 | (pre 0-10um) / (10D10S 11-20um) | 0.014* | 0.003 | 0.067 |
| (10D10S 31-40um) / (pre 21-30um) | 0.151* | 0.085 | 0.267 | (pre 0-10um) / (pre 11-20um) | 0.007* | 0.001 | 0.032 |
| (pre 31-40um) / (10D10S 0-10um) | 5.343* | 1.416 | 20.159 | (pre 0-10um) / (10D10S 21-30um) | 0.002* | 0.001 | 0.011 |
| (pre 31-40um) / (pre 0-10um) | 23.3* | 4.813 | 112.796 | (pre 0-10um) / (pre 21-30um) | 0.003* | 0.001 | 0.015 |
| (pre 31-40um) / (10D10S 11-20um) | 0.338* | 0.183 | 0.621 | (pre 0-10um) / (10D10S 31-40um) | 0.02* | 0.004 | 0.095 |
| (pre 31-40um) / (pre 11-20um) | 0.157* | 0.079 | 0.311 | (pre 0-10um) / (pre 31-40um) | 0.043* | 0.009 | 0.208 |
| (pre 31-40um) / (10D10S 21-30um) | 0.057* | 0.031 | 0.103 | (pre 0-10um) / (10D10S 41-50um) | 0.079* | 0.016 | 0.384 |
| (pre 31-40um) / (pre 21-30um) | 0.072* | 0.036 | 0.143 | (pre 0-10um) / (pre 41-50um) | 0.24 | 0.045 | 1.265 |
| (pre 31-40um) / (10D10S 31-40um) | 0.478* | 0.257 | 0.887 | (pre 0-10um) / (10D10S 51-60um) | 0.123* | 0.021 | 0.713 |
| (10D10S 41-50um) / (10D10S 0-10um) | 2.916 | 0.765 | 11.107 | (pre 0-10um) / (pre 51-60um) | 0.333 | 0.061 | 1.83 |
| (10D10S 41-50um) / (pre 0-10um) | 12.715* | 2.605 | 62.055 | (pre 0-10um) / (10D10S 61-70um) | 0.209 | 0.025 | 1.744 |
| (10D10S 41-50um) / (10D10S 11-20um) | 0.184* | 0.098 | 0.346 | (pre 0-10um) / (pre 61-70um) | 0.914 | 0.133 | 6.267 |
| (10D10S 41-50um) / (pre 11-20um) | 0.085* | 0.042 | 0.173 | (pre 0-10um) / (10D10S 71-80um) | 0.284 | 0.021 | 3.785 |
| (10D10S 41-50um) / (10D10S 21-30um) | 0.031* | 0.017 | 0.058 | (pre 0-10um) / (pre 71-80um) | 1.183 | 0.09 | 15.621 |
| (10D10S 41-50um) / (pre 21-30um) | 0.039* | 0.019 | 0.08 | (pre 0-10um) / (10D10S 81-90um) | 0.199 | 0.015 | 2.665 |
| (10D10S 41-50um) / (10D10S 31-40um) | 0.261* | 0.137 | 0.494 | (pre 0-10um) / (10D10S 101-110um) | 0.185 | 0.014 | 2.47 |
| (10D10S 41-50um) / (pre 31-40um) | 0.546 | 0.26 | 1.147 | (pre 0-10um) / (pre 101-110um) | 0.824 | 0.062 | 10.896 |
| (pre 41-50um) / (10D10S 0-10um) | 0.957 | 0.229 | 4.001 | (pre 0-10um) / (pre 141-150um) | 1.183 | 0.09 | 15.621 |
| (pre 41-50um) / (pre 0-10um) | 4.175 | 0.791 | 22.049 | (10D10S 11-20um) / (pre 11-20um) | 0.464* | 0.264 | 0.815 |
| (pre 41-50um) / (10D10S 11-20um) | 0.06* | 0.027 | 0.136 | (10D10S 11-20um) / (10D10S 21-30um) | 0.168* | 0.107 | 0.264 |
| (pre 41-50um) / (pre 11-20um) | 0.028* | 0.012 | 0.067 | (10D10S 11-20um) / (pre 21-30um) | 0.214* | 0.122 | 0.374 |
| (pre 41-50um) / (10D10S 21-30um) | 0.01* | 0.005 | 0.023 | (10D10S 11-20um) / (10D10S 31-40um) | 1.415 | 0.877 | 2.281 |
| (pre 41-50um) / (pre 21-30um) | 0.013* | 0.005 | 0.031 | (10D10S 11-20um) / (pre 31-40um) | 2.963* | 1.61 | 5.451 |
| (pre 41-50um) / (10D10S 31-40um) | 0.086* | 0.038 | 0.193 | (10D10S 11-20um) / (10D10S 41-50um) | 5.429* | 2.891 | 10.198 |
| (pre 41-50um) / (pre 31-40um) | 0.179* | 0.073 | 0.44 | (10D10S 11-20um) / (pre 41-50um) | 16.534* | 7.368 | 37.105 |
| (pre 41-50um) / (10D10S 41-50um) | 0.328* | 0.132 | 0.818 | (10D10S 11-20um) / (10D10S 51-60um) | 8.502* | 3.179 | 22.738 |
| (10D10S 51-60um) / (10D10S 0-10um) | 1.862 | 0.401 | 8.651 | (10D10S 11-20um) / (pre 51-60um) | 22.999* | 9.477 | 55.816 |
| (10D10S 51-60um) / (pre 0-10um) | 8.12* | 1.403 | 47.007 | (10D10S 11-20um) / (10D10S 61-70um) | 14.454* | 3.093 | 67.54 |
| (10D10S 51-60um) / (10D10S 11-20um) | 0.118* | 0.044 | 0.315 | (10D10S 11-20um) / (pre 61-70um) | 63.063* | 17.852 | 222.771 |
| (10D10S 51-60um) / (pre 11-20um) | 0.055* | 0.019 | 0.153 | (10D10S 11-20um) / (10D10S 71-80um) | 19.636* | 2.308 | 167.043 |
| (10D10S 51-60um) / (10D10S 21-30um) | 0.02* | 0.007 | 0.053 | (10D10S 11-20um) / (pre 71-80um) | 81.688* | 9.696 | 688.244 |
| (10D10S 51-60um) / (pre 21-30um) | 0.025* | 0.009 | 0.071 | (10D10S 11-20um) / (10D10S 81-90um) | 13.762* | 1.609 | 117.71 |
| (10D10S 51-60um) / (10D10S 31-40um) | 0.166* | 0.062 | 0.448 | (10D10S 11-20um) / (10D10S 101-110um) | 12.741* | 1.487 | 109.134 |
| (10D10S 51-60um) / (pre 31-40um) | 0.348 | 0.121 | 1.005 | (10D10S 11-20um) / (pre 101-110um) | 56.917* | 6.747 | 480.174 |
| (10D10S 51-60um) / (10D10S 41-50um) | 0.639 | 0.219 | 1.865 | (10D10S 11-20um) / (pre 141-150um) | 81.688* | 9.696 | 688.236 |
| (10D10S 51-60um) / (pre 41-50um) | 1.945 | 0.594 | 6.362 | (pre 11-20um) / (10D10S 21-30um) | 0.363* | 0.21 | 0.629 |
| (pre 51-60um) / (10D10S 0-10um) | 0.688 | 0.157 | 3.011 | (pre 11-20um) / (pre 21-30um) | 0.46* | 0.242 | 0.877 |
| (pre 51-60um) / (pre 0-10um) | 3.002 | 0.546 | 16.488 | (pre 11-20um) / (10D10S 31-40um) | 3.05* | 1.719 | 5.413 |
| (pre 51-60um) / (10D10S 11-20um) | 0.043* | 0.018 | 0.106 | (pre 11-20um) / (pre 31-40um) | 6.387* | 3.211 | 12.707 |
| (pre 51-60um) / (pre 11-20um) | 0.02* | 0.008 | 0.052 | (pre 11-20um) / (10D10S 41-50um) | 11.705* | 5.778 | 23.715 |
| (pre 51-60um) / (10D10S 21-30um) | 0.007* | 0.003 | 0.018 | (pre 11-20um) / (pre 41-50um) | 35.646* | 14.957 | 84.953 |
| (pre 51-60um) / (pre 21-30um) | 0.009* | 0.004 | 0.024 | (pre 11-20um) / (10D10S 51-60um) | 18.329* | 6.517 | 51.552 |
| (pre 51-60um) / (10D10S 31-40um) | 0.062* | 0.025 | 0.15 | (pre 11-20um) / (pre 51-60um) | 49.585* | 19.335 | 127.158 |
| (pre 51-60um) / (pre 31-40um) | 0.129* | 0.049 | 0.34 | (pre 11-20um) / (10D10S 61-70um) | 31.162* | 6.456 | 150.412 |
| (pre 51-60um) / (10D10S 41-50um) | 0.236* | 0.088 | 0.631 | (pre 11-20um) / (pre 61-70um) | 135.96* | 36.997 | 499.637 |
| (pre 51-60um) / (pre 41-50um) | 0.719 | 0.238 | 2.172 | (pre 11-20um) / (10D10S 71-80um) | 42.333* | 4.86 | 368.716 |
| (pre 51-60um) / (10D10S 51-60um) | 0.37 | 0.107 | 1.277 | (pre 11-20um) / (pre 71-80um) | 176.115* | 20.414 | 1519.333 |
| (10D10S 61-70um) / (10D10S 0-10um) | 1.095 | 0.157 | 7.631 | (pre 11-20um) / (10D10S 81-90um) | 29.671* | 3.388 | 259.806 |
| (10D10S 61-70um) / (pre 0-10um) | 4.776 | 0.574 | 39.773 | (pre 11-20um) / (10D10S 101-110um) | 27.469* | 3.133 | 240.873 |
| (10D10S 61-70um) / (10D10S 11-20um) | 0.069* | 0.015 | 0.323 | (pre 11-20um) / (pre 101-110um) | 122.71* | 14.205 | 1059.992 |
| (10D10S 61-70um) / (pre 11-20um) | 0.032* | 0.007 | 0.155 | (pre 11-20um) / (pre 141-150um) | 176.113* | 20.414 | 1519.313 |
| (10D10S 61-70um) / (10D10S 21-30um) | 0.012* | 0.003 | 0.054 | (10D10S 21-30um) / (pre 21-30um) | 1.268 | 0.735 | 2.189 |
| (10D10S 61-70um) / (pre 21-30um) | 0.015* | 0.003 | 0.071 | (10D10S 21-30um) / (10D10S 31-40um) | 8.402* | 5.298 | 13.323 |
| (10D10S 61-70um) / (10D10S 31-40um) | 0.098* | 0.021 | 0.459 | (10D10S 21-30um) / (pre 31-40um) | 17.595* | 9.679 | 31.984 |
| (10D10S 61-70um) / (pre 31-40um) | 0.205 | 0.042 | 1.006 | (10D10S 21-30um) / (10D10S 41-50um) | 32.243* | 17.373 | 59.842 |
| (10D10S 61-70um) / (10D10S 41-50um) | 0.376 | 0.076 | 1.859 | (10D10S 21-30um) / (pre 41-50um) | 98.191* | 44.18 | 218.231 |
| (10D10S 61-70um) / (pre 41-50um) | 1.144 | 0.214 | 6.122 | (10D10S 21-30um) / (10D10S 51-60um) | 50.489* | 19.016 | 134.051 |
| (10D10S 61-70um) / (10D10S 51-60um) | 0.588 | 0.1 | 3.448 | (10D10S 21-30um) / (pre 51-60um) | 136.586* | 56.778 | 328.57 |
| (10D10S 61-70um) / (pre 51-60um) | 1.591 | 0.286 | 8.855 | (10D10S 21-30um) / (10D10S 61-70um) | 85.839* | 18.461 | 399.134 |
| (pre 61-70um) / (10D10S 0-10um) | 0.251 | 0.045 | 1.412 | (10D10S 21-30um) / (pre 61-70um) | 374.515* | 106.643 | 1315.239 |
| (pre 61-70um) / (pre 0-10um) | 1.095 | 0.16 | 7.509 | (10D10S 21-30um) / (10D10S 71-80um) | 116.611* | 13.754 | 988.638 |
| (pre 61-70um) / (10D10S 11-20um) | 0.016* | 0.004 | 0.056 | (10D10S 21-30um) / (pre 71-80um) | 485.123* | 57.777 | 4073.308 |
| (pre 61-70um) / (pre 11-20um) | 0.007* | 0.002 | 0.027 | (10D10S 21-30um) / (10D10S 81-90um) | 81.73* | 9.588 | 696.665 |
| (pre 61-70um) / (10D10S 21-30um) | 0.003* | 0.001 | 0.009 | (10D10S 21-30um) / (10D10S 101-110um) | 75.665* | 8.864 | 645.906 |
| (pre 61-70um) / (pre 21-30um) | 0.003* | 0.001 | 0.012 | (10D10S 21-30um) / (pre 101-110um) | 338.015* | 40.204 | 2841.869 |
| (pre 61-70um) / (10D10S 31-40um) | 0.022* | 0.006 | 0.08 | (10D10S 21-30um) / (pre 141-150um) | 485.12* | 57.777 | 4073.256 |
| (pre 61-70um) / (pre 31-40um) | 0.047* | 0.013 | 0.176 | (pre 21-30um) / (10D10S 31-40um) | 6.624* | 3.743 | 11.722 |
| (pre 61-70um) / (10D10S 41-50um) | 0.086* | 0.023 | 0.326 | (pre 21-30um) / (pre 31-40um) | 13.871* | 6.987 | 27.539 |
| (pre 61-70um) / (pre 41-50um) | 0.262 | 0.063 | 1.09 | (pre 21-30um) / (10D10S 41-50um) | 25.42* | 12.573 | 51.394 |
| (pre 61-70um) / (10D10S 51-60um) | 0.135* | 0.029 | 0.623 | (pre 21-30um) / (pre 41-50um) | 77.411* | 32.54 | 184.156 |
| (pre 61-70um) / (pre 51-60um) | 0.365 | 0.084 | 1.587 | (pre 21-30um) / (10D10S 51-60um) | 39.804* | 14.171 | 111.803 |
| (pre 61-70um) / (10D10S 61-70um) | 0.229 | 0.033 | 1.591 | (pre 21-30um) / (pre 51-60um) | 107.68* | 42.059 | 275.687 |
| (10D10S 71-80um) / (10D10S 0-10um) | 0.806 | 0.07 | 9.29 | (pre 21-30um) / (10D10S 61-70um) | 67.673* | 14.034 | 326.332 |
| (10D10S 71-80um) / (pre 0-10um) | 3.516 | 0.264 | 46.782 | (pre 21-30um) / (pre 61-70um) | 295.256* | 80.433 | 1083.834 |
| (10D10S 71-80um) / (10D10S 11-20um) | 0.051* | 0.006 | 0.433 | (pre 21-30um) / (10D10S 71-80um) | 91.933* | 10.562 | 800.194 |
| (10D10S 71-80um) / (pre 11-20um) | 0.024* | 0.003 | 0.206 | (pre 21-30um) / (pre 71-80um) | 382.457* | 44.362 | 3297.28 |
| (10D10S 71-80um) / (10D10S 21-30um) | 0.009* | 0.001 | 0.073 | (pre 21-30um) / (10D10S 81-90um) | 64.434* | 7.363 | 563.836 |
| (10D10S 71-80um) / (pre 21-30um) | 0.011* | 0.001 | 0.095 | (pre 21-30um) / (10D10S 101-110um) | 59.652* | 6.807 | 522.746 |
| (10D10S 71-80um) / (10D10S 31-40um) | 0.072* | 0.008 | 0.615 | (pre 21-30um) / (pre 101-110um) | 266.481* | 30.869 | 2300.412 |
| (10D10S 71-80um) / (pre 31-40um) | 0.151 | 0.017 | 1.33 | (pre 21-30um) / (pre 141-150um) | 382.455* | 44.362 | 3297.238 |
| (10D10S 71-80um) / (10D10S 41-50um) | 0.277 | 0.031 | 2.453 | (10D10S 31-40um) / (pre 31-40um) | 2.094* | 1.128 | 3.889 |
| (10D10S 71-80um) / (pre 41-50um) | 0.842 | 0.09 | 7.914 | (10D10S 31-40um) / (10D10S 41-50um) | 3.838* | 2.025 | 7.274 |
| (10D10S 71-80um) / (10D10S 51-60um) | 0.433 | 0.043 | 4.36 | (10D10S 31-40um) / (pre 41-50um) | 11.687* | 5.171 | 26.412 |
| (10D10S 71-80um) / (pre 51-60um) | 1.171 | 0.121 | 11.338 | (10D10S 31-40um) / (10D10S 51-60um) | 6.009* | 2.234 | 16.168 |
| (10D10S 71-80um) / (10D10S 61-70um) | 0.736 | 0.055 | 9.88 | (10D10S 31-40um) / (pre 51-60um) | 16.257* | 6.656 | 39.706 |
| (10D10S 71-80um) / (pre 61-70um) | 3.212 | 0.28 | 36.891 | (10D10S 31-40um) / (10D10S 61-70um) | 10.217* | 2.178 | 47.918 |
| (pre 71-80um) / (10D10S 0-10um) | 0.194 | 0.017 | 2.214 | (10D10S 31-40um) / (pre 61-70um) | 44.576* | 12.561 | 158.187 |
| (pre 71-80um) / (pre 0-10um) | 0.845 | 0.064 | 11.156 | (10D10S 31-40um) / (10D10S 71-80um) | 13.879* | 1.627 | 118.395 |
| (pre 71-80um) / (10D10S 11-20um) | 0.012* | 0.001 | 0.103 | (10D10S 31-40um) / (pre 71-80um) | 57.741* | 6.835 | 487.81 |
| (pre 71-80um) / (pre 11-20um) | 0.006* | 0.001 | 0.049 | (10D10S 31-40um) / (10D10S 81-90um) | 9.728* | 1.134 | 83.428 |
| (pre 71-80um) / (10D10S 21-30um) | 0.002* | 0 | 0.017 | (10D10S 31-40um) / (10D10S 101-110um) | 9.006* | 1.049 | 77.349 |
| (pre 71-80um) / (pre 21-30um) | 0.003* | 0 | 0.023 | (10D10S 31-40um) / (pre 101-110um) | 40.232* | 4.756 | 340.335 |
| (pre 71-80um) / (10D10S 31-40um) | 0.017* | 0.002 | 0.146 | (10D10S 31-40um) / (pre 141-150um) | 57.741* | 6.835 | 487.804 |
| (pre 71-80um) / (pre 31-40um) | 0.036* | 0.004 | 0.317 | (pre 31-40um) / (10D10S 41-50um) | 1.833 | 0.872 | 3.853 |
| (pre 71-80um) / (10D10S 41-50um) | 0.066* | 0.008 | 0.584 | (pre 31-40um) / (pre 41-50um) | 5.581* | 2.271 | 13.715 |
| (pre 71-80um) / (pre 41-50um) | 0.202 | 0.022 | 1.885 | (pre 31-40um) / (10D10S 51-60um) | 2.87 | 0.995 | 8.279 |
| (pre 71-80um) / (10D10S 51-60um) | 0.104 | 0.01 | 1.039 | (pre 31-40um) / (pre 51-60um) | 7.763* | 2.942 | 20.481 |
| (pre 71-80um) / (pre 51-60um) | 0.282 | 0.029 | 2.701 | (pre 31-40um) / (10D10S 61-70um) | 4.879 | 0.994 | 23.953 |
| (pre 71-80um) / (10D10S 61-70um) | 0.177 | 0.013 | 2.356 | (pre 31-40um) / (pre 61-70um) | 21.285* | 5.675 | 79.836 |
| (pre 71-80um) / (pre 61-70um) | 0.772 | 0.068 | 8.793 | (pre 31-40um) / (10D10S 71-80um) | 6.628 | 0.752 | 58.441 |
| (pre 71-80um) / (10D10S 71-80um) | 0.24 | 0.012 | 4.755 | (pre 31-40um) / (pre 71-80um) | 27.572* | 3.157 | 240.822 |
| (10D10S 81-90um) / (10D10S 0-10um) | 1.15 | 0.099 | 13.318 | (pre 31-40um) / (10D10S 81-90um) | 4.645 | 0.524 | 41.178 |
| (10D10S 81-90um) / (pre 0-10um) | 5.016 | 0.375 | 67.046 | (pre 31-40um) / (10D10S 101-110um) | 4.3 | 0.484 | 38.176 |
| (10D10S 81-90um) / (10D10S 11-20um) | 0.073* | 0.008 | 0.621 | (pre 31-40um) / (pre 101-110um) | 19.211* | 2.197 | 168.013 |
| (10D10S 81-90um) / (pre 11-20um) | 0.034* | 0.004 | 0.295 | (pre 31-40um) / (pre 141-150um) | 27.572* | 3.157 | 240.819 |
| (10D10S 81-90um) / (10D10S 21-30um) | 0.012* | 0.001 | 0.104 | (10D10S 41-50um) / (pre 41-50um) | 3.045* | 1.222 | 7.59 |
| (10D10S 81-90um) / (pre 21-30um) | 0.016* | 0.002 | 0.136 | (10D10S 41-50um) / (10D10S 51-60um) | 1.566 | 0.536 | 4.573 |
| (10D10S 81-90um) / (10D10S 31-40um) | 0.103* | 0.012 | 0.882 | (10D10S 41-50um) / (pre 51-60um) | 4.236* | 1.585 | 11.323 |
| (10D10S 81-90um) / (pre 31-40um) | 0.215 | 0.024 | 1.908 | (10D10S 41-50um) / (10D10S 61-70um) | 2.662 | 0.538 | 13.176 |
| (10D10S 81-90um) / (10D10S 41-50um) | 0.395 | 0.044 | 3.518 | (10D10S 41-50um) / (pre 61-70um) | 11.615* | 3.067 | 43.991 |
| (10D10S 81-90um) / (pre 41-50um) | 1.201 | 0.127 | 11.35 | (10D10S 41-50um) / (10D10S 71-80um) | 3.617 | 0.408 | 32.08 |
| (10D10S 81-90um) / (10D10S 51-60um) | 0.618 | 0.061 | 6.252 | (10D10S 41-50um) / (pre 71-80um) | 15.046* | 1.712 | 132.197 |
| (10D10S 81-90um) / (pre 51-60um) | 1.671 | 0.172 | 16.259 | (10D10S 41-50um) / (10D10S 81-90um) | 2.535 | 0.284 | 22.603 |
| (10D10S 81-90um) / (10D10S 61-70um) | 1.05 | 0.078 | 14.159 | (10D10S 41-50um) / (10D10S 101-110um) | 2.347 | 0.263 | 20.956 |
| (10D10S 81-90um) / (pre 61-70um) | 4.582 | 0.397 | 52.885 | (10D10S 41-50um) / (pre 101-110um) | 10.483* | 1.192 | 92.229 |
| (10D10S 81-90um) / (10D10S 71-80um) | 1.427 | 0.071 | 28.53 | (10D10S 41-50um) / (pre 141-150um) | 15.046* | 1.712 | 132.196 |
| (10D10S 81-90um) / (pre 71-80um) | 5.936 | 0.299 | 117.875 | (pre 41-50um) / (10D10S 51-60um) | 0.514 | 0.157 | 1.682 |
| (10D10S 101-110um) / (10D10S 0-10um) | 1.242 | 0.107 | 14.403 | (pre 41-50um) / (pre 51-60um) | 1.391 | 0.46 | 4.203 |
| (10D10S 101-110um) / (pre 0-10um) | 5.418 | 0.405 | 72.507 | (pre 41-50um) / (10D10S 61-70um) | 0.874 | 0.163 | 4.678 |
| (10D10S 101-110um) / (10D10S 11-20um) | 0.078* | 0.009 | 0.672 | (pre 41-50um) / (pre 61-70um) | 3.814 | 0.918 | 15.852 |
| (10D10S 101-110um) / (pre 11-20um) | 0.036* | 0.004 | 0.319 | (pre 41-50um) / (10D10S 71-80um) | 1.188 | 0.126 | 11.162 |
| (10D10S 101-110um) / (10D10S 21-30um) | 0.013* | 0.002 | 0.113 | (pre 41-50um) / (pre 71-80um) | 4.941 | 0.531 | 46.01 |
| (10D10S 101-110um) / (pre 21-30um) | 0.017* | 0.002 | 0.147 | (pre 41-50um) / (10D10S 81-90um) | 0.832 | 0.088 | 7.864 |
| (10D10S 101-110um) / (10D10S 31-40um) | 0.111* | 0.013 | 0.954 | (pre 41-50um) / (10D10S 101-110um) | 0.771 | 0.081 | 7.29 |
| (10D10S 101-110um) / (pre 31-40um) | 0.233 | 0.026 | 2.064 | (pre 41-50um) / (pre 101-110um) | 3.442 | 0.369 | 32.098 |
| (10D10S 101-110um) / (10D10S 41-50um) | 0.426 | 0.048 | 3.805 | (pre 41-50um) / (pre 141-150um) | 4.941 | 0.531 | 46.01 |
| (10D10S 101-110um) / (pre 41-50um) | 1.298 | 0.137 | 12.277 | (10D10S 51-60um) / (pre 51-60um) | 2.705 | 0.783 | 9.348 |
| (10D10S 101-110um) / (10D10S 51-60um) | 0.667 | 0.066 | 6.763 | (10D10S 51-60um) / (10D10S 61-70um) | 1.7 | 0.29 | 9.968 |
| (10D10S 101-110um) / (pre 51-60um) | 1.805 | 0.185 | 17.586 | (10D10S 51-60um) / (pre 61-70um) | 7.418* | 1.605 | 34.288 |
| (10D10S 101-110um) / (10D10S 61-70um) | 1.134 | 0.084 | 15.313 | (10D10S 51-60um) / (10D10S 71-80um) | 2.31 | 0.229 | 23.26 |
| (10D10S 101-110um) / (pre 61-70um) | 4.95 | 0.428 | 57.197 | (10D10S 51-60um) / (pre 71-80um) | 9.609 | 0.963 | 95.903 |
| (10D10S 101-110um) / (10D10S 71-80um) | 1.541 | 0.077 | 30.849 | (10D10S 51-60um) / (10D10S 81-90um) | 1.619 | 0.16 | 16.384 |
| (10D10S 101-110um) / (pre 71-80um) | 6.411 | 0.323 | 127.456 | (10D10S 51-60um) / (10D10S 101-110um) | 1.499 | 0.148 | 15.189 |
| (10D10S 101-110um) / (10D10S 81-90um) | 1.08 | 0.054 | 21.705 | (10D10S 51-60um) / (pre 101-110um) | 6.695 | 0.67 | 66.903 |
| (pre 101-110um) / (10D10S 0-10um) | 0.278 | 0.024 | 3.182 | (10D10S 51-60um) / (pre 141-150um) | 9.608 | 0.963 | 95.902 |
| (pre 101-110um) / (pre 0-10um) | 1.213 | 0.092 | 16.028 | (pre 51-60um) / (10D10S 61-70um) | 0.628 | 0.113 | 3.497 |
| (pre 101-110um) / (10D10S 11-20um) | 0.018* | 0.002 | 0.148 | (pre 51-60um) / (pre 61-70um) | 2.742 | 0.63 | 11.93 |
| (pre 101-110um) / (pre 11-20um) | 0.008* | 0.001 | 0.07 | (pre 51-60um) / (10D10S 71-80um) | 0.854 | 0.088 | 8.264 |
| (pre 101-110um) / (10D10S 21-30um) | 0.003* | 0 | 0.025 | (pre 51-60um) / (pre 71-80um) | 3.552 | 0.37 | 34.068 |
| (pre 101-110um) / (pre 21-30um) | 0.004* | 0 | 0.032 | (pre 51-60um) / (10D10S 81-90um) | 0.598 | 0.062 | 5.822 |
| (pre 101-110um) / (10D10S 31-40um) | 0.025* | 0.003 | 0.21 | (pre 51-60um) / (10D10S 101-110um) | 0.554 | 0.057 | 5.397 |
| (pre 101-110um) / (pre 31-40um) | 0.052* | 0.006 | 0.455 | (pre 51-60um) / (pre 101-110um) | 2.475 | 0.258 | 23.767 |
| (pre 101-110um) / (10D10S 41-50um) | 0.095* | 0.011 | 0.839 | (pre 51-60um) / (pre 141-150um) | 3.552 | 0.37 | 34.067 |
| (pre 101-110um) / (pre 41-50um) | 0.29 | 0.031 | 2.709 | (10D10S 61-70um) / (pre 61-70um) | 4.363 | 0.629 | 30.277 |
| (pre 101-110um) / (10D10S 51-60um) | 0.149 | 0.015 | 1.493 | (10D10S 61-70um) / (10D10S 71-80um) | 1.358 | 0.101 | 18.233 |
| (pre 101-110um) / (pre 51-60um) | 0.404 | 0.042 | 3.881 | (10D10S 61-70um) / (pre 71-80um) | 5.652 | 0.424 | 75.251 |
| (pre 101-110um) / (10D10S 61-70um) | 0.254 | 0.019 | 3.385 | (10D10S 61-70um) / (10D10S 81-90um) | 0.952 | 0.071 | 12.836 |
| (pre 101-110um) / (pre 61-70um) | 1.108 | 0.097 | 12.634 | (10D10S 61-70um) / (10D10S 101-110um) | 0.881 | 0.065 | 11.898 |
| (pre 101-110um) / (10D10S 71-80um) | 0.345 | 0.017 | 6.831 | (10D10S 61-70um) / (pre 101-110um) | 3.938 | 0.295 | 52.489 |
| (pre 101-110um) / (pre 71-80um) | 1.435 | 0.073 | 28.222 | (10D10S 61-70um) / (pre 141-150um) | 5.651 | 0.424 | 75.251 |
| (pre 101-110um) / (10D10S 81-90um) | 0.242 | 0.012 | 4.806 | (pre 61-70um) / (10D10S 71-80um) | 0.311 | 0.027 | 3.577 |
| (pre 101-110um) / (10D10S 101-110um) | 0.224 | 0.011 | 4.454 | (pre 61-70um) / (pre 71-80um) | 1.295 | 0.114 | 14.754 |
| (pre 141-150um) / (10D10S 0-10um) | 0.194 | 0.017 | 2.214 | (pre 61-70um) / (10D10S 81-90um) | 0.218 | 0.019 | 2.519 |
| (pre 141-150um) / (pre 0-10um) | 0.845 | 0.064 | 11.156 | (pre 61-70um) / (10D10S 101-110um) | 0.202 | 0.017 | 2.335 |
| (pre 141-150um) / (10D10S 11-20um) | 0.012* | 0.001 | 0.103 | (pre 61-70um) / (pre 101-110um) | 0.903 | 0.079 | 10.292 |
| (pre 141-150um) / (pre 11-20um) | 0.006* | 0.001 | 0.049 | (pre 61-70um) / (pre 141-150um) | 1.295 | 0.114 | 14.753 |
| (pre 141-150um) / (10D10S 21-30um) | 0.002* | 0 | 0.017 | (10D10S 71-80um) / (pre 71-80um) | 4.16 | 0.21 | 82.297 |
| (pre 141-150um) / (pre 21-30um) | 0.003* | 0 | 0.023 | (10D10S 71-80um) / (10D10S 81-90um) | 0.701 | 0.035 | 14.015 |
| (pre 141-150um) / (10D10S 31-40um) | 0.017* | 0.002 | 0.146 | (10D10S 71-80um) / (10D10S 101-110um) | 0.649 | 0.032 | 12.989 |
| (pre 141-150um) / (pre 31-40um) | 0.036* | 0.004 | 0.317 | (10D10S 71-80um) / (pre 101-110um) | 2.899 | 0.146 | 57.395 |
| (pre 141-150um) / (10D10S 41-50um) | 0.066* | 0.008 | 0.584 | (10D10S 71-80um) / (pre 141-150um) | 4.16 | 0.21 | 82.296 |
| (pre 141-150um) / (pre 41-50um) | 0.202 | 0.022 | 1.885 | (pre 71-80um) / (10D10S 81-90um) | 0.168 | 0.008 | 3.346 |
| (pre 141-150um) / (10D10S 51-60um) | 0.104 | 0.01 | 1.039 | (pre 71-80um) / (10D10S 101-110um) | 0.156 | 0.008 | 3.101 |
| (pre 141-150um) / (pre 51-60um) | 0.282 | 0.029 | 2.701 | (pre 71-80um) / (pre 101-110um) | 0.697 | 0.035 | 13.701 |
| (pre 141-150um) / (10D10S 61-70um) | 0.177 | 0.013 | 2.356 | (pre 71-80um) / (pre 141-150um) | 1 | 0.051 | 19.645 |
| (pre 141-150um) / (pre 61-70um) | 0.772 | 0.068 | 8.793 | (10D10S 81-90um) / (10D10S 101-110um) | 0.926 | 0.046 | 18.603 |
| (pre 141-150um) / (10D10S 71-80um) | 0.24 | 0.012 | 4.755 | (10D10S 81-90um) / (pre 101-110um) | 4.136 | 0.208 | 82.208 |
| (pre 141-150um) / (pre 71-80um) | 1 | 0.051 | 19.646 | (10D10S 81-90um) / (pre 141-150um) | 5.936 | 0.299 | 117.874 |
| (pre 141-150um) / (10D10S 81-90um) | 0.168 | 0.008 | 3.346 | (10D10S 101-110um) / (pre 101-110um) | 4.467 | 0.225 | 88.89 |
| (pre 141-150um) / (10D10S 101-110um) | 0.156 | 0.008 | 3.101 | (10D10S 101-110um) / (pre 141-150um) | 6.411 | 0.323 | 127.455 |
| (pre 141-150um) / (pre 101-110um) | 0.697 | 0.035 | 13.701 | (pre 101-110um) / (pre 141-150um) | 1.435 | 0.073 | 28.222 |

^1^Lower confidence limit, ^2^Upper confidence limit, * Significant pairwise comparisons
